# Essential role of the amino-terminal region of Drosha for the Microprocessor function

**DOI:** 10.1101/2022.10.12.509557

**Authors:** Amit Prabhakar, Song Hu, Jin Tang, Prajakta Ghatpande, Giorgio Lagna, Xuan Jiang, Akiko Hata

**Author notes:** Corresponding Authors Xuan Jiang, PhD, Molecular Cancer Research Center, Sun Yat-Sen University School of Medicine, Xiuqi Building Room 216 No. 132, Waihuan East Rd., Guangzhou University City, Panyu District, Guangzhou, Guangdong, 511400, P.R. China, /Fax, Akiko Hata, PhD (Lead Contact), Cardiovascular Research Institute, University of California, San Francisco, Mail Box: 3118, 555 Mission Bay Blvd. South, SCVRB Room 252T, San Francisco, CA 94143, USA, /.

## Abstract

The ribonuclease (RNase) III enzyme Drosha enables processing of microRNA (miRNA) primary transcripts (pri-miRNAs) and control of ribosomal protein (RP) biogenesis by the Microprocessor. The extensively studied carboxyl-terminal half of Drosha is sufficient to crop pri-miRNAs in vitro, but the function of the evolutionarily conserved Drosha amino-terminal region (NTR) is unknown, despite it harboring mutations linked to a hereditary vascular disorder. Here, we provide evidence that, when replacing the endogenous Drosha, a mutant missing the NTR (ΔN-Drosha) fails to associate with endogenous pri-miRNAs and to support the expression of all miRNAs, except the miR-183 cluster. Surprisingly, Argonaute2 (Ago2) associates with and cleaves pri-miR-183 in ΔN-Drosha cells and in Drosha-depleted cells. ΔN-Drosha is also unable to inhibit RP biogenesis upon serum starvation. Thus, Drosha NTR is essential for pri-miRNA processing and RP biogenesis in vivo, and Ago2 can process the miR-183 cluster in the absence of Drosha.

## Introduction

Drosha and DiGeorge syndrome critical region 8 (Dgcr8, also known as Pasha) are components of the Microprocessor, a complex responsible for the biogenesis of miRNAs (Kim et al., 2009; Lee et al., 2007). The RNase III enzyme Drosha processes long pri-miRNAs to generate precursor-miRNAs (pre-miRNAs) with short stem-loop (hairpin) structures in the nucleus (Kim *et al*., 2009; Lee *et al*., 2007), which then undergo secondary processing by Dicer in the cytoplasm to generate small RNA duplexes of ∼22-nucleotides (nt)(Ha and Kim, 2014). The RNA duplexes are then loaded onto an Argonaute (Ago) protein to form an RNA-induced silencing complex (RISC), which unwind RNA duplexes, remove the passenger strands, and induce destabilization and translational repression of target mRNAs through recognition of the sequence that is partially complementary to the miRNA (Ha and Kim, 2014). The Microprocessor-mediated processing is regulated by physiological stimuli, for example, upon activation of the TGF--family of growth factor signaling (Hata and Lieberman, 2015). All four human Ago proteins (Ago1-4) incorporate miRNAs in RISC, but Ago2 is distinct because it is capable of miRNA-directed target RNA cleavage through the intrinsic RNA slicing activity (Okamura et al., 2004). It is also found that Ago2, instead of Dicer, processes pre-miR-451(Kretov et al., 2020; Yang and Lai, 2010; Yang et al., 2010), suggesting that the slicing activity of Ago2 has broader functions beyond miRNA-mediated mRNA silencing.

In the carboxyl (C)-terminal region of Drosha [hereafter referred to as CTR; amino acid (aa) 875-1374], two RNase III domains (aa 875-1056 and aa 1107-1233) and one double-stranded RNA-binding domain (dsRBD) (aa 1260-1344) are essential for the processing of pri-miRNAs(Kim *et al*., 2009). Conversely, the functions of the conserved proline (P)-rich region (aa 69-164) and arginine/serine (RS)-rich region (aa 217-315) in the amino (N)-terminal part of Drosha (hereafter referred to as NTR) remain to be elucidated(Kim *et al*., 2009). Since a truncated version of Drosha missing the NTR (aa 390-1365) is sufficient to interact with Dgcr8 and process pri-miRNAs in vitro, the Drosha-NTR was initially considered dispensable for Drosha function as a component of the Microprocessor (Han et al., 2004), hence it has been incompletely studied.

Several disease-associated alleles of Drosha have been identified in humans, including loss-of-function mutations in the RNase III domains of Drosha in patients with the pediatric renal cancer Wilms tumor (Kruber et al., 2018; Rakheja et al., 2014; Spreafico et al., 2016; Torrezan et al., 2014; Walz et al., 2015; Wegert et al., 2015) and missense mutations in the P-rich and the RS-rich regions of Drosha (*e*.*g*., P100L and R279L) in patients with the familial vascular disorder hereditary hemorrhagic telangiectasia (HHT)(Hata and Lagna, 2019; Jiang et al., 2018). While Wilms tumor mutations in the Drosha CTR affect the RNase III activity (Kruber *et al*., 2018; Rakheja *et al*., 2014; Spreafico *et al*., 2016; Torrezan *et al*., 2014; Walz *et al*., 2015; Wegert *et al*., 2015), the effect of HHT mutations in the Drosha NTR are yet-to-be explored. We demonstrated an important role of the Drosha NTR in the control of ribosomal protein (RP) synthesis upon nutrients deprivation (Jiang et al., 2021b). Briefly, the Microprocessor complex associates with the 5’-oligopyrimidine (5’TOP) tract of nascent RP gene (RPG) transcripts and facilitates RNA polymerase II (RNAPII) elongation via the RNA helicase Ddx5, an auxiliary component of the Microprocessor (Jiang *et al*., 2021b). Upon nutrient starvation, Drosha is degraded by the proteasome-dependent mechanism, which results in repression of RP synthesis, reduction of ribosomes, and inhibition of global protein synthesis (Jiang *et al*., 2021b). Here we expand the analysis of Drosha NTR by generating cell lines expressing a Drosha-NTR truncation mutant (ΔN-Drosha) from the endogenous locus. We observe that the Drosha NTR is essential for pri-miRNA processing, its control by TGF-, and the regulation of RP synthesis. ΔN-Drosha localizes to the nucleus and interacts with Dgcr8, but it is defective in pri-miRNA processing and leads to globally diminished miRNA levels except for the miR-183/96/182 cluster. We found that the processing of pri-miR-183/96/182 (hereafter referred to as pri-miR-183) is mediated by Ago2. Finally, ΔN-Drosha is resistant to the ubiquitin-dependent degradation, and therefore, ΔN-Drosha cells do not repress RP synthesis, global translation, and cell proliferation in response to nutrient deprivation. Our study sheds light on the essential role of Drosha-NTR in the canonical and noncanonical functions of the Microprocessor.

## Results

### Establishing cell lines expressing Drosha without the NTR

The NTR of Drosha contains evolutionarily conserved P-rich and R/S-rich regions with unclear functions (**Fig. 1a**). To study the function of the NTR, we planned to generate by reverse genetics Drosha mutants lacking this region. In the human *Drosha* gene, exon 5 (ex5) encodes aa 7-284, which comprise the P-rich region (aa 69-164) and most of the R/S-rich region (aa 217-315). To generate a cell line in which ex5 of *Drosha* is deleted, we transfected two guide RNAs (gRNA1 and gRNA2) that target the intron upstream and downstream of ex5, respectively, together with the plasmid encoding the Cas9 enzyme (**Supplementary Fig. S1**). We isolated 10 stable clones, examined the status of ex5 by PCR analysis, and identified the heterozygous clone 7 (with one *Drosha* allele lacking ex5; Δex5/+) and the homozygous mutant clone 8 (with both *Drosha* alleles lacking ex5; Δex5/Δex5) (**Supplementary Fig. S1 and S2**). RNA-seq data detected no reads (in clone 8) and a smaller number of reads corresponding to ex5 in clone 7 (**Fig. 1b, orange box**).

**Fig. 1.**
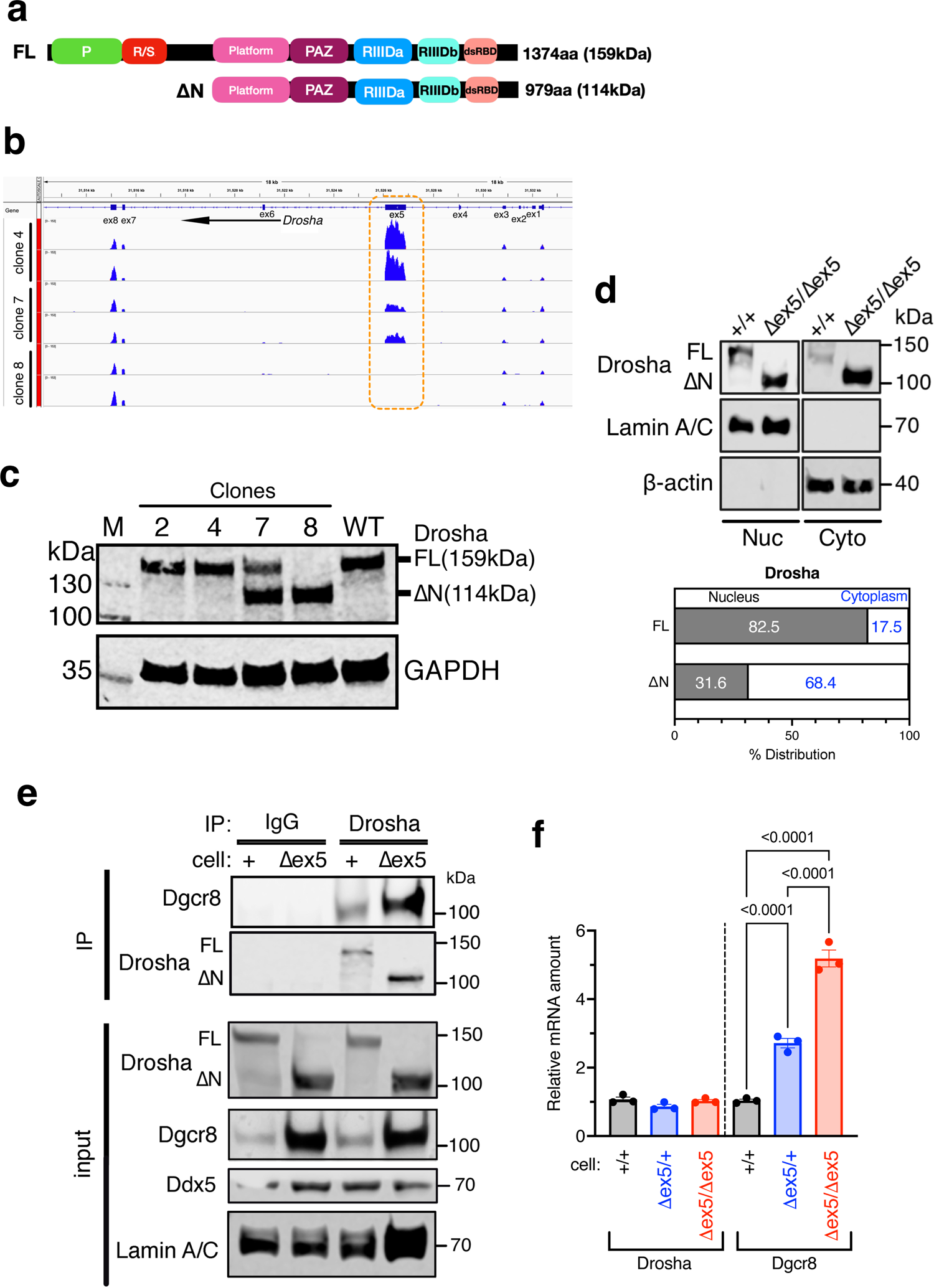
Generation of cell lines expressing Drosha truncated in the NTR. **a**. Schematic diagram of the domain structure of Drosha FL and ΔN-Drosha protein and the human Drosha gene structure (light blue). P: Pro-rich region, R/S: Arg/Ser-rich region, PAZ: Piwi Argonaut and Zwille domain, RIIID: RNase III domain, dsRBD: double-stranded RNA binding domain. **b**. RNA-seq data of clones 4, 7, and 8 corresponding to the ex1-8 of the Drosha gene are shown. Each clone was sequenced twice. **c**. Total cell lysates of clones 2, 4, 7, 8 or the original HEK293T cells (WT) were subjected to immunoblot analysis of Drosha and GAPDH (loading control). **d**. Nuclear (Nuc) and cytoplasmic (Cyto) fraction of +/+ cells (FL) and Δex5/Δex5 cells (ΔN) cells were subjected to immunoblot analysis of Drosha, Lamin A/C (control for the Nuc fraction), and β-actin (control for the Cyto fraction) (top). Relative distribution (%) of FL and ΔN-Drosha in the nucleus vs cytoplasm is shown (bottom). **e**. Co-immunoprecipitation of Drosha (FL or ΔN-Drosha) and Dgcr8 in nuclear extracts of +/+ cells (+) and Δex5/Δex5 cells (Δex5). As control, non-specific IgG (control) was applied. The amount of Drosha after IP is shown. Input samples were subjected to immunoblot analyses of Drosha, Dgcr8, Ddx5, and Lamin A/C (loading control). **f**. The level of the Drosha and Dgcr8 mRNA relative to GAPDH mRNA in +/+ cells (black), Δex5/+ cells (blue), and Δex5/Δex5 cells (red) were measured by qRT-PCR and plotted as mean ± SEM. n=3 independent experiments.

Clones 2 and 4 retained both *Drosha* wild type alleles (+/+) (**Supplementary Fig. S1 and S2**). The translation of the Δex5 allele in clones 7 and 8 is predicted to start at methionine-396 to generate a mutant Drosha lacking the NTR (ΔN-Drosha; aa 396-1374) with a molecular weight (M.W.) of 114 kDa. Immunoblot analysis confirmed that clones 2 and 4 (+/+) express a full length (FL; aa 1-1374) Drosha (FL) with a M.W. of 159 kDa identical to the wild-type protein in the original HEK293T cells (WT) (**Fig. 1c**) while clone 8 (Δex5/Δex5) expresses only ΔN-Drosha (**Fig. 1c**). Clone 7 (Δex5/+) expressed both FL and ΔN-Drosha (**Fig. 1c**), confirming its heterozygosity. To examine the subcellular localization of FL and ΔN-Drosha, the nuclear and cytoplasmic fractions of +/+ and Δex5/Δex5 cells were subjected to immunoblot analysis with anti-Drosha antibodies. FL in +/+ cells was found predominantly in the nucleus while ΔN-Drosha in Δex5/Δex5 cells was predominantly in the cytoplasm (**Fig. 1d**). We interpret this result as due to the loss of a nuclear localization signal (NLS) in the RS-rich region in ΔN-Drosha(Lee et al., 2006). Immunoprecipitation of Drosha followed by immunoblot with an anti-Dgcr8 antibody indicated that both FL and ΔN-Drosha were able to interact with Dgcr8 (**Fig. 1e**). We noticed that ΔN-Drosha protein was 2.5-fold more abundant than FL, despite being expressed from the same locus (**Fig. 1e**). Furthermore, Dgcr8 protein in Δex5/Δex5 cells was 8-fold more abundant than in +/+ cells (**Fig. 1e, input**). qRT-PCR analysis showed that Drosha mRNA in +/+, Δex5/+, and Δex5/Δex5 cells was similarly expressed (**Fig. 1f, Drosha**), indicating that the higher amount of ΔN-Drosha protein in Δex5/Δex5 cells is likely due to increased protein stability. Unlike Drosha mRNA, the amount of Dgcr8 mRNA in Δex5/Δex5 cells was 5-fold higher than in +/+ cells (**Fig. 1f, Dgcr8 and Supplementary Fig. S3**). Considering that Dgcr8 mRNA is cleaved by Drosha and degraded(Han et al., 2009), the increased level of Dgcr8 mRNA in Δex5/Δex5 cells may suggest a defective catalytic activity of ΔN-Drosha.

### Depletion of global miRNAs in ΔN-Drosha cells

To compare the pri-miRNA processing activity of wild-type and mutant Drosha, we compared FL and ΔN-Drosha from +/+ and Δex5/Δex5 cells in an in vitro processing (IVP) assay performed using a fluorescein-conjugated substrate (pri-let-7b) (**Fig. 2a, top left**). The catalytic activity of ΔN-Drosha was ∼80% lower than that of FL (**Fig. 2a, top right**). This is consistent with the increased amount of the Dgcr8 mRNA in ΔN-Drosha cells (**Fig. 1f, input and Supplementary Fig. S3**). When global miRNA expression was examined in +/+, Δex5/+, and Δex5/Δex5 cells by small RNA-seq, 98% of miRNAs (598 out of 611) were diminished in Δex5/Δex5 cells compared to +/+ cells, indicating an essential, general role for Drosha NTR in pri-miRNA processing (**Fig. 2b, left**). However, specific miRNAs, including miR-96-5p, miR-129-1-3p, miR-182-5p, miR-183-5p, miR-1247-3p, and miR-4775, were more expressed in Δex5/Δex5 cells than +/+ cells (**Fig. 2b, left, blue**). There were no significant changes in miRNA levels between Δex5/+ and +/+ cells (**Fig. 2b, right**). Among the miRNAs that did not diminish in Δex5/Δex5 cells were miR-103a1-5p, miR-25-3p, miR-224-5p, miR-93-5p, miR26b-5p, miR-26a1-5p, miR139-5p, and miR-30c-5p (**Fig. 2b, left, orange**), which are known “intronic-miRNAs” whose processing is Drosha-independent(Lin et al., 2006). qRT-PCR analysis validated the small RNA-seq result, confirming that most miRNAs were greatly reduced in Δex5/Δex5 cells (**Fig. 2c, black vs red**) and less reduced in Δex5/+ cells (**Fig. 2c, black vs blue**). The amount of Drosha-independent miR-103a was similar in all 3 clones (**Fig. 2c**). The RNA-immunoprecipitation (RIP) assay showed that the association of ΔN-Drosha in Δex5/Δex5 cells with the hairpin of pri-let-7b or pri-miR-21, compared to FL in +/+ cells, was reduced by 70% and 99%, respectively (**Fig. 2d**), suggesting that the NTR significantly contributes to the ability of Drosha to stably associate with and crop pri-miRNAs in the Microprocessor.

**Fig. 2.**
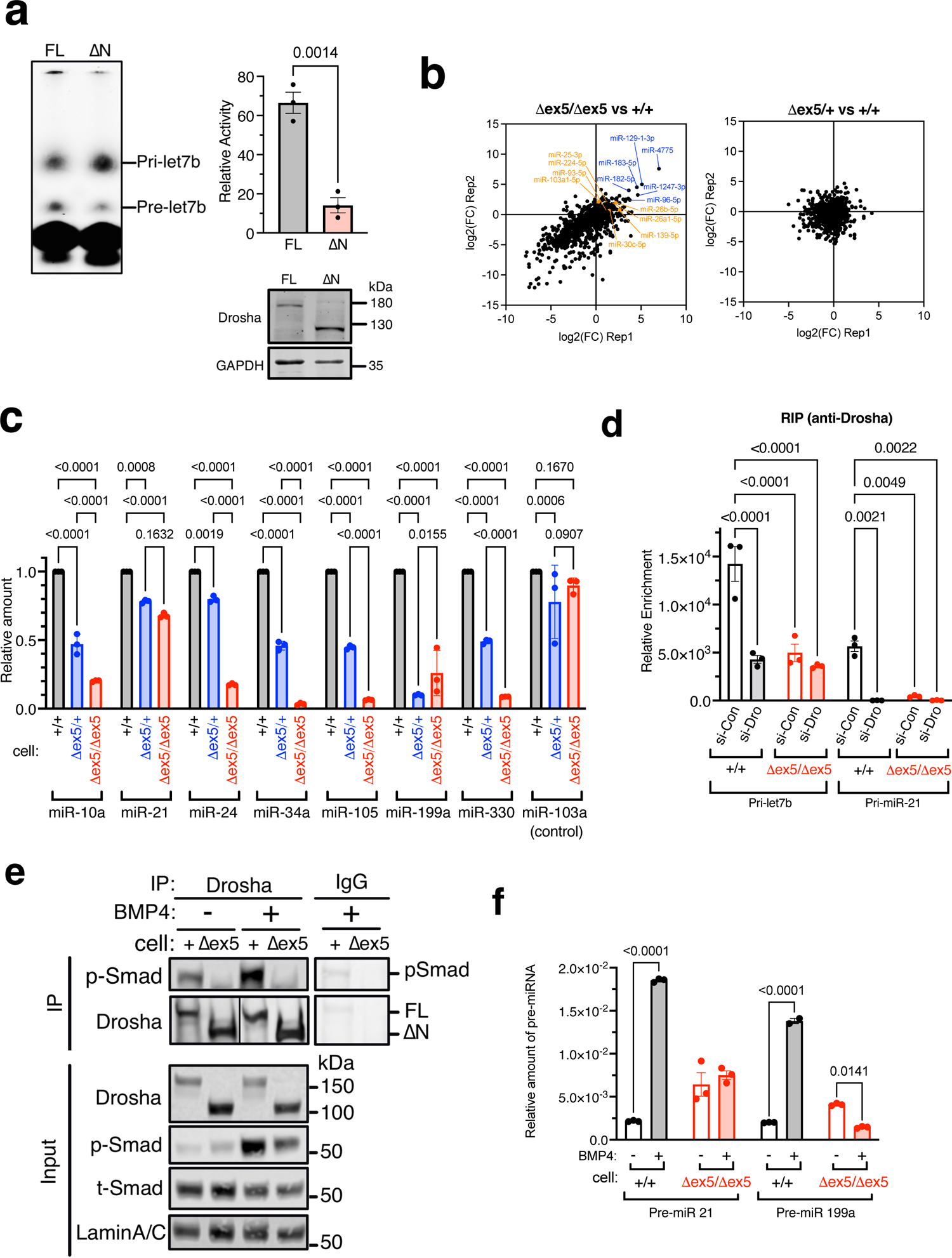
The NTR of Drosha is essential for the processing of pri-miRNAs. **a**. In vitro processing of pri-let-7b. A fluorescein-labeled substrate (pri-let-7b) was incubated with FL and ΔN-Drosha from +/+ cells (clone 4) and Δex5/Δex5 (clone 8) cells, respectively. Immunoblot analysis of Drosha indicates the amount of Drosha-FL and ΔN-Drosha added to the processing reaction (bottom). After the reaction, the processing product (pre-let7b) was separated from the substrate by PAGE (top left) and the processing activity was quantitated, normalized by Drosha amount, and shown (top right). **b**. Small RNA-seq analysis of miRNAs in +/+ cells (clone 4), Δex5/+ cells (clone 7), and Δex5/Δex5 cells (clone 8). Two libraries from two independent samples were generated from each clone. The proportion of miRNA reads in the small RNA sequencing libraries from +/+ cells and Δex5/Δex5 cells (left) or +/+ cells and Δex5/+ cells(right) are compared and the fold change (FC) of miRNAs (log_2_FC) are shown in scatter plot. miRNAs shown in orange are intronic miRNA whose processing is independent of Drosha. **c**. qRT-PCR analysis showing the amount of miR-10, −21, −24, −34a, −103a1, − 105, −199a, and −330 relative to U6 snRNA in +/+ cells (black), Δex5/+ cells (blue), and Δex5/Δex5 cells (red). miR-103a1 is an intronic miRNA. The result was plotted as means ± SEM. n=3. **d**. Association of Drosha with pri-let7b or pri-miR-21 was assessed by RIP assay in +/+ cells (black) and Δex5/Δex5 cells (red). The number of pri-miRNAs in the anti-Drosha antibody immunoprecipitates (IP) was quantitated by qRT-PCR in triplicates. Relative enrichment of IP over input was plotted as mean± SEM. **e**. Co-immunoprecipitation of Drosha (FL or ΔN) and phosphorylated-Smad1/5/8 (p-Smad) was examined in +/+ cells (FL) or Δex5/Δex5 cells (ΔN-Drosha) with or without 1 nM BMP4 treatment for 2 hr. As control, non-specific IgG (control) was applied. The amount of Drosha after IP is also shown. Input samples were subjected to immunoblot analyses of Drosha, p-Smad, total-Smad1 (t-Smad), and Lamin A/C (loading control). **f**. qRT-PCR analysis of pre-miR-21 and pre-miR-199a in +/+ cells (FL) or Δex5/Δex5 cells (ΔN-Drosha) cells with or without 1 nM BMP4 stimulation for 2 hr. The result is plotted as mean± SEM. n=3 independent experiments.

The Microprocessor activity is regulated by different proteins that associate with the Microprocessor, such as the Smads(Blahna and Hata, 2013; Hata and Lieberman, 2015). Smads 1, 5, and 8, the signal transducers of the bone morphogenetic proteins (BMPs), interact with Drosha upon BMP4 stimulation and induce the processing of specific miRNAs, such as miR-21 and miR-199a(Davis et al., 2008). IP-immunoblot analyses showed that Smad1/5/8 proteins phosphorylated by the BMP receptor kinase upon BMP4 stimulation (p-Smad) associated with Drosha-FL in +/+ cells but not with ΔN-Drosha in Δex5/Δex5 cells (**Fig. 2e, IP**), although p-Smads were detected in the input samples of Δex5/Δex5 cells stimulated with BMP4 (**Fig. 2f, Input**). In +/+ cells, the level of pre-miR-21 and pre-miR-199a increased 9-fold and 7-fold upon BMP4 stimulation, respectively (**Fig. 2f, black bars**), but neither pre-miR-21 nor pre-miR-199a increased upon BMP4 treatment of Δex5/Δex5 cells (**Fig. 2f, red bars**). These results indicate the NTR of Drosha is essential for the BMP4-dependent rise in Microprocessor activity through association with Smad proteins.

### Increased production of RPs in cells expressing ΔN-Drosha

The Microprocessor potentiates the transcription of ∼80 ribosomal protein (RP) genes (RPGs) by binding to the 5’TOP sequence shared by all RPGs(Jiang *et al*., 2021b). A ChIP assay indicated that both FL and ΔN-Drosha associate with the *Rps2, Rps10*, and *Rpl28* gene loci at similar levels (**Fig. 3a**), indicating that Drosha NTR is dispensable for the interaction with RPGs. Our previous studies showed that, upon serum starvation, Drosha translocates from the nucleus to the cytoplasm and is degraded after Nedd4-dependent ubiquitination, resulting in the repression of RPG transcription(Jiang *et al*., 2021b). We repeated these experiments in cells harboring wild-type or mutant Drosha. Upon serum starvation (1% serum) for 16 h in +/+ cells, ∼90% of FL translocated to the cytoplasm (**Fig. 3b**); FL and RPs diminished in a time-dependent manner (**Fig. 3c**); and the amount of RPG mRNAs also rapidly declined (**Supplementary Fig. S4**), as expected from our previous results. However, in Δex5/Δex5 cells after serum starvation, more than 50% of ΔN-Drosha remained in the nucleus (**Fig. 3b**) and the amount of ΔN-Drosha and RPs remained higher than in +/+ cells (**Fig. 3c**). Consistently, the amount of Gata1 protein (whose translation is sensitive to the change in ribosome abundance(Jiang *et al*., 2021b)) was rapidly reduced to ∼20% in +/+ cells 6 h after serum starvation, but decreased only 3% in ΔN-Drosha cells, in accordance with the higher amount of RPs in these cells (**Fig. 3c**). An in vitro puromycin incorporation assay(Aviner et al., 2013) showed that the amount of puromycin-labeled nascent proteins was reduced to 38% in +/+ cells after serum starvation for 16 h (**Fig. 3d**), indicating a reduction of global translation. Conversely, there was no significant reduction of translation in ΔN-Drosha-expressing cells (**Fig. 3d**), which is consistent with the persistence of RP biogenesis in these cells upon serum starvation (**Fig. 3c**). When the proliferation of +/+ and Δex5/Δex5 cells was compared in normal growth media (10% serum) and starvation media (1% serum), the doubling time (*Td*) of +/+ cells increased from 20 h (10% serum) to 56 h (1% serum) (**Fig. 3e, black lines**), indicating a slower proliferation in 1% serum. Conversely, cells expressing ΔN-Drosha proliferated at a similar rate in 10% serum (*Td*=21 hr) and 1% serum (*Td*=21 hr) media (**Fig. 3e, red lines**). These results demonstrate that the NTR of Drosha is dispensable for the interaction with RPGs, but necessary for nutrient deprivation-mediated degradation of Drosha. As a result, ΔN-Drosha cells are unable to adapt to the change in the availability of nutrients and cannot modulate ribosome abundance and cell growth rate.

**Fig. 3.**
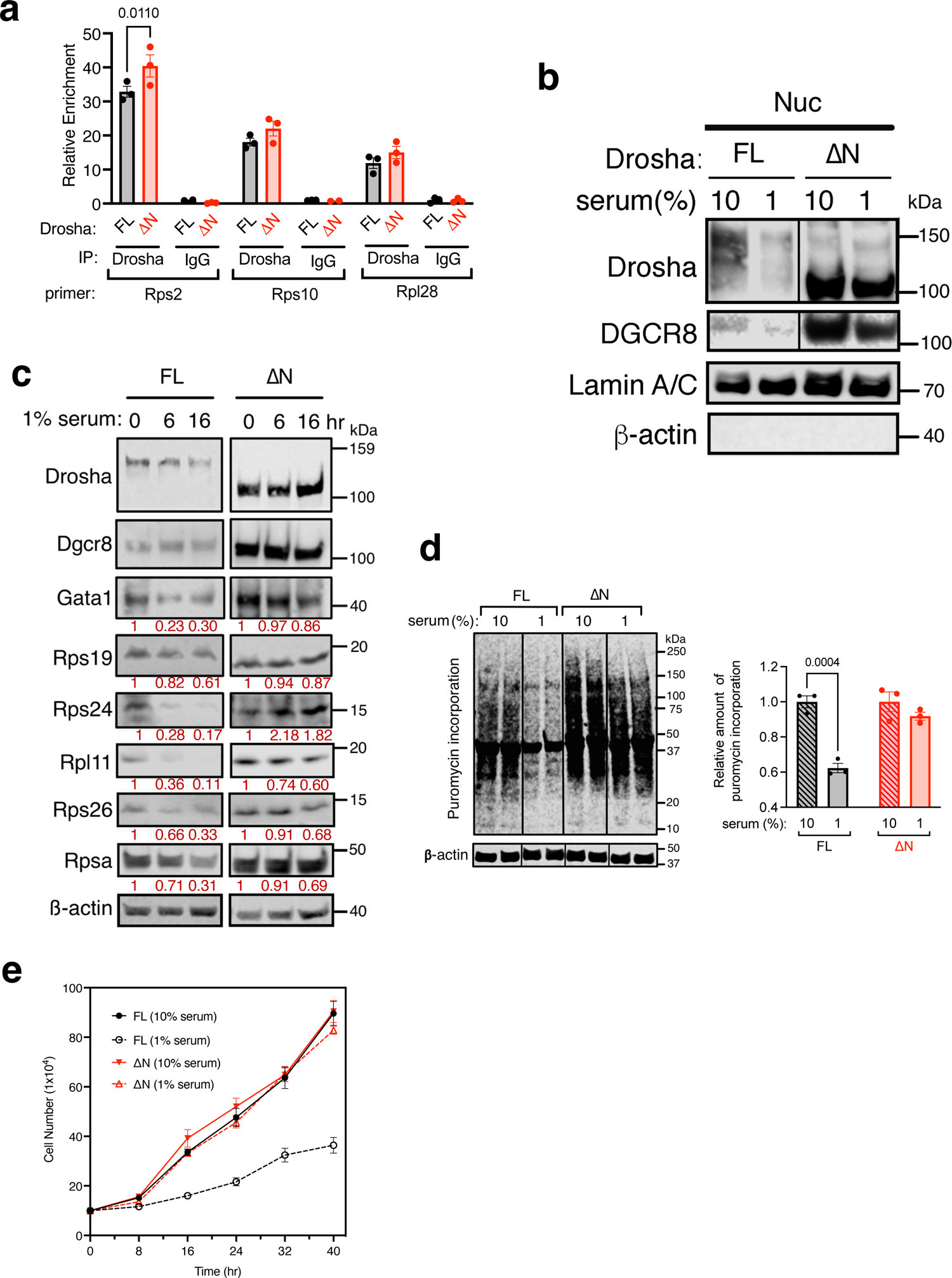
Maintenance of ribosomal protein abundance, protein synthesis, and growth in ΔN-Drosha cells under serum starvation. **a**. Association of Drosha with the *Rps2, Rps10*, and *Rpl28* transcripts. ChIP-qPCR analysis was performed by using anti-Drosha antibody (Drosha) or nonspecific IgG (IgG; negative control) in FL or ΔN-Drosha cells. The result is shown as a fold enrichment over input (Mean ± SEM). n=3 independent experiments. **b**. FL or ΔN-Drosha cells cultured in 10% serum or 1% serum for 16 hr, followed by the preparation of nuclear (Nuc) lysates and subjected to immunoblot for Drosha, Dgcr8, Lamin A/C (loading control for the Nuc fraction) and β-actin (loading control for the cytoplasmic fraction). **c**. FL or ΔN-Drosha cells were serum starved (1% serum) for 0, 6, or 16 hr, followed by immunoblot analysis for Drosha, Dgcr8, Gata1, RPs (Rps19, Rps24, Rpl11, Rps26, Rpsa), and β-actin (loading control). Red numbers indicate the relative amount of the protein shown above during serum starvation. **d**. Equal number of FL or ΔN-Drosha cells were cultured in normal growth media (10% serum) or serum starvation media (1% serum) for 16 hr, followed by puromycin treatment for 10 min and immunoblot analysis with anti-puromycin antibody and anti--actin antibody (loading control) (left). The relative abundance of puromycin-incorporated proteins normalized by β-actin is shown (right) as mean± SEM. n=3 per group. Unpaired two-tail t-test was used for the statistical analysis. **e**. FL or ΔN-Drosha cells were cultured in growth media (10% serum) or starvation media (1% serum) and the cell number was counted at 8, 16, 24, 32, and 40 hr after the media change. The result is plotted as mean± SEM. n=5 independent samples.

### Drosha-independent, Ago2-dependent processing of the miR-183 cluster

Among the miRNAs expressed in ΔN-Drosha (Δex5/Δex5) cells at a level equivalent or slightly higher than FL (+/+) cells were miR-183, −96, and −182 (**Fig. 2b, left and Supplementary Fig. S5**). These three miRNAs belong to the miR-183 cluster and are transcribed as a single long polycistronic transcript (pri-miR-183) with three hairpin structures corresponding to mature miR-183, −96, and −182 (Dambal et al., 2015). A qRT-PCR analysis confirmed that these three miRNAs were expressed at similar or slightly higher amounts in ΔN-Drosha cells compared to FL cells (**Fig. 4a**). The pri-miR-183 transcript was similar in FL and ΔN-Drosha cells (**Fig. 4a**), indicating no change in the miR-183 cluster gene transcription. When Drosha was depleted by siRNA (si-Dro) in FL cells, the levels of miR-183 cluster miRNAs were slightly elevated (**Fig. 4b**) while other miRNAs (miR-21, −24, −105, and −330) were diminished (**Fig. 4b and Supplementary Fig. S6**). Next, we conducted an IVP assay, in which fluorescein-labeled pri-miR-183 or pri-let-7b were incubated with Drosha from control siRNA-transfected (si-Con) or si-Drosha-transfected (si-Dro) HEK293T cells. The result showed that depletion of Drosha (si-Dro) represses the processing of pri-let-7b, but not of pri-miR-183 (**Fig. 4c**), further confirming that pri-miR-183 processing is Drosha-independent. The small RNA-seq data revealed a large fraction of miR-183 and miR-96 sequence variants (isomiRs) (**Table 1, red**), which contain 1 or 2 additional nucleotides at the 5’-end with respect to the reference sequence (**Table 1, blue**), in ΔN-Drosha cells. We speculate that the cleavage of these pri-miRNAs in ΔN-Drosha cells occurred 1- or 2-nt upstream of the conventional Drosha cleavage site. Together, these results support the hypothesis that an enzyme other than Drosha is processing pri-miR-183 when Drosha is depleted or inactive.

**Table 1:**
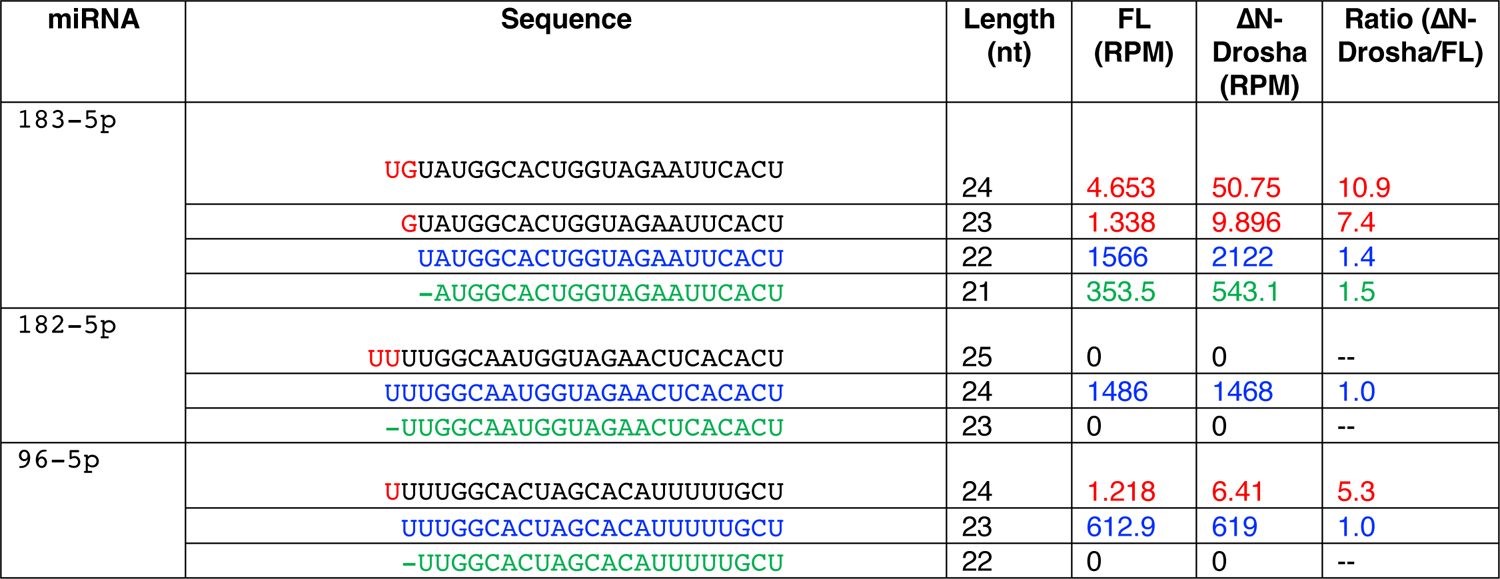
IsomiRs of miR-183 and miR-96 are found in ΔN-Drosha cells. The average number reads (reads per million; RPM) of miR-183-5p, −182-5p, and −96-5p in FL or ΔN-Drosha cells are shown. Blue sequences are reference sequences of miR-183-5p, −182-5p, −96-5p by miRBase. Additional nucleotides at the 5’-end in respect of the reference sequence are shown in red. The ratio (ΔN-Drosha/FL) of the isomiR-183 that is 1-nt shorter than the reference sequence (green) is close to the ratio of total number of miR-183-5p reads. The ratio of isomiR-183-5p (10.9 and 7.4) and isomiR-95-5p (5.3) are higher than the ratio of total number of miR-183-5p (1.4) and miR-96-5p (1.0), respectively.

A high-throughput sequencing following cross-linking immunoprecipitation (HITS-CLIP) analysis showed that miR-183/96/182 are efficiently bound by Ago2 (Chu-Tan et al., 2021). Furthermore, Ago2 is uniquely capable of directly cleaving highly complementary RNA targets(Liu et al., 2004; Meister et al., 2004; Okamura *et al*., 2004). We found that ∼40% of Ago2 is localized in the nucleus in FL and ΔN-Drosha cells (**Supplementary Fig. S7**). Thus, we hypothesized that nuclear Ago2 might play a role in pri-miR-183 processing when Drosha expression or activity is compromised. As expected, an association of Drosha-FL with both pri-miR-183 and pri-miR-21 was detected in FL cells by RIP assay (**Fig. 4d, black, si-Con**). This interaction was diminished by Drosha depletion (**Fig. 4d, black, si-Dro**), but unchanged by Ago2 depletion (**Fig. 4d, black, si-Ag2**). Unlike in FL cells, no association of Drosha with pri-miR-183 or pri-miR-21 was detected in ΔN-Drosha cells (**Fig. 4d, red, si-Con**), although the amounts of pri-miR-183 and pri-miR-21 were similar in ΔN-Drosha and FL cells (**Supplementary Fig. S8A**). Instead of interacting with Drosha, pri-miR-183 associated with Ago2 in ΔN-Drosha cells (**Fig. 4e, red, si-Con**). When Ago2 was depleted by siRNA (si-Ag2), the RIP signal detected in ΔN-Drosha cells (si-Con) was abolished (**Fig. 4e, red, si-Ag2**), indicating that the signal is due to the interaction between Ago2 and pri-miR-183. When Drosha was depleted in ΔN-Drosha cells, the Ago2-pri-miR-183 interaction was dampened (**Fig. 4e, red, si-Dro**), suggesting that ΔN-Drosha facilitates the Ago2-pri-miR-183 interaction, likely through the association with Dgcr8. Despite similar amounts of Ago2 protein being immunoprecipitated by anti-Ago2 antibody from ΔN-Drosha and FL cells (**Supplementary Fig. S8B**), the Ago2-pri-miR183 interaction was not detected in FL cells (**Fig. 4e, black**). Ago2 did not associate with pri-miR-21 in ΔN-Drosha or FL cells (**Fig. 4e**), indicating the specificity of the Ago2-pri-miR-183 interaction. In FL cells, Ago2 did not associate with pri-miR-183 (**Fig. 4e, black**), but associate with it when Drosha was depleted by siRNA (si-Dro) (**Fig. 4e, black**). These results suggest that Ago2-dependent processing of pri-miR-183 occurs when Drosha is absent or defective. The knock-down of Ago2 by siRNA (si-Ag2) in FL cells did not affect the amount of miR-183 cluster miRNAs (miR-183, −96, and −182) (**Fig. 4f, black**). In contrast, Ago2 knock-down reduced the amount of 3 miRNAs in the miR-183 cluster, but not miR-21 or intronic miR-103a, in ΔN-Drosha cells (**Fig. 4f, red**), indicating that Ago2 plays a role in the pri-miR-183 processing in ΔN-Drosha cells. RIP assay with anti-Dgcr8 antibody confirmed that pri-miR-183 was bound by Dgcr8 in ΔN-Drosha cells (**Fig. 4g**), but when Ago2 was depleted, the amount of pri-miR-183 bound by Dgcr8 diminished (**Fig. 4g**). Unlike pri-miR-183, Dgcr8 association with the pri-miR-21 was similar in the presence or absence of Ago2 (**Fig. 4g**). We detected the interaction between Dgcr8 and Ago2 in both FL and ΔN-Drosha cells by IP-immunoblot analysis (**Supplementary Fig. S9, orange rectangle**). Therefore, we speculate that the association of Ago2 and Dgcr8 with the pri-miR-183 is facilitated through the complex formation between Ago2 and Dgcr8. Together, these data demonstrate that pri-miR-183 can be processed by Ago2 and Dgcr8 when Drosha is absent or catalytically impaired (**Fig. 4h**). To our knowledge, this is the first example of Drosha-independent Ago2-dependent cleavage of pri-miRNA.

**Fig. 4.**
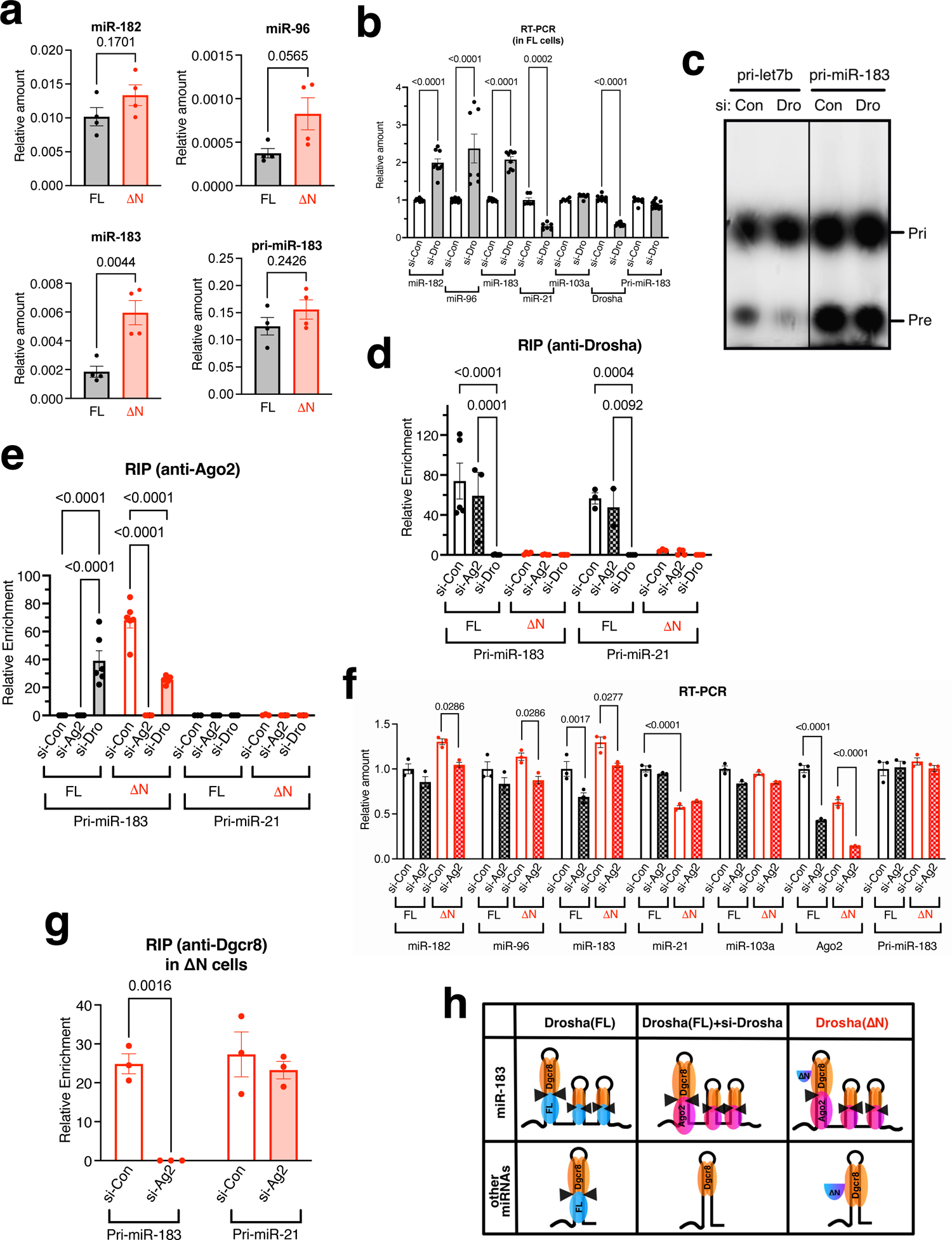
The processing of miR-183 cluster is independent of Drosha. **a**. qRT-PCR analysis of miR-182, −96, and −183 (normalized to U6 snRNA), and pri-miR-183 (normalized to GAPDH) in FL and ΔN-Drosha cells. The result is plotted as mean± SEM. n=4 independent experiments. **b**. FL cells transfected with siRNA against non-specific control (si-Con) or Drosha (si-Dro) were subjected to qRT-PCR analysis of miR-182, −96, −183, control miRNAs (miR-21, and −103a) (normalized to U6 snRNA), Drosha mRNA, and pri-miR-183 (normalized to GAPDH) was performed in triplicates. The result is plotted as mean± SEM. n=2-3 independent experiments. **c**. In vitro processing assay of pri-let-7b and pri-miR-183. A fluorescein-labeled substrate (pri-let-7b or pri-miR-183) was incubated with Drosha isolated from HEK293T cells transfected with non-specific control siRNA (si-Con) or si-Drosha (si-Dro), followed by the separation of the processing product (pre-let-7b or pre-miR-183) by PAGE. **d**. RIP assay to access the interaction between Drosha (FL or ΔN) and pri-miR-183 or pri-miR-21 in FL and ΔN-Drosha cells transfected with siRNA against non-specific control (si-Con), Ago2 (si-Ag2) or Drosha (si-Dro). The amount of pri-miR-183 or pri-miR21 in the immunoprecipitates of anti-Drosha antibody or non-specific IgG (control) was quantitated by qRT-PCR in triplicates. Relative enrichment of anti-Drosha antibody IP over IgG control is plotted as mean± SEM. n=2-3 independent experiments. **e**. Association of Ago2 with pri-miR-183 or pri-miR-21 is assessed by RIP assay in FL and ΔN-Drosha cells transfected with non-specific control siRNA (si-Con), Ago2 (si-Ag2), or Drosha (si-Dro). The amount of pri-miR-183 or pri-miR-21 in the immunoprecipitates of anti-Ago2 antibody or non-specific IgG (control) was quantitated by qRT-PCR in triplicates. Relative enrichment of anti-Drosha antibody IP over IgG control is plotted as mean± SEM. n=2-3 independent experiments. **f**. qRT-PCR analysis of miR-182, −96, 183, −21, and −103a, Ago2, and pri-miR-183/96/182 in FL or ΔN-Drosha cells transfected with siRNA against non-specific control siRNA (si-Con) or Ago2 (si-Ag2). The amount of miRNAs and Drosha mRNA normalized to U6 snRNA and Drosha mRNAs normalized to GAPDH is plotted as mean± SEM. n=3 independent experiments. **g**. ΔN-Drosha cells were transfected with non-specific control siRNA (si-Con) or si-Ag2, followed by RIP assay with IP with anti-Dgcr8 antibody or non-specific IgG (control). The amount of pri-miR-183 or pri-miR-21 in the immunoprecipitates was quantitated by qRT-PCR. Relative enrichment of IP (anti-Dgcr8 antibody) over input was plotted as mean± SEM. n=3. **h**. A schematic representation of Drosha-independent, Ago2-dependent processing of the pri-miR-183. In FL cells, the pri-miR-183 is processed by Drosha and Dgcr8 like other pri-miRNAs. When Drosha-FL is depleted, Ago2 and Dgcr8 process the pri-miR-183 in FL cells. In ΔN-Drosha cells, the processing of pri-miRNAs is blunted, however, the pri-miR-183 is processed by Ago2 and Dgcr8.

## Discussion

In this work, we demonstrated that Drosha-ΔN fails to associate with pri-miRNAs, and thus the Drosha NTR is essential for the Microprocessor activity. A previous in vitro study on a N-terminus truncated Drosha (ΔN-390; aa 390-1374), also lacking the P-rich region and the R/S-rich region as the ΔN-Drosha used in this study, showed that processing of pri-let7a1 by ΔN-390 was comparable to that of wild type Drosha (WT) (Han *et al*., 2004), leading to the conclusion that the Drosha-NTR (aa 1-389) was dispensable for the catalytic activity. Based on the immunoblot presented, though, it is possible that a higher amount of ΔN-390 than the WT protein might have been used in the IVP assay, decreasing the difference in activity between the two Drosha forms(Han *et al*., 2004).

Cryo-electron microscopy structure of the NTR-truncated Drosha (aa 353-1372) and a partial Dgcr8 (amino acid 223-751) with a pri-miRNA-16-2 demonstrate that the basal tip of the central domain (CED; amino acid 353-960) of Drosha wraps around the dsRNA-single stranded RNA (ssRNA) junction of the pri-miRNA(Partin et al., 2020). Because ΔN-Drosha lacks the N-terminus 43 aa out of 608 aa of the CED, it is plausible that ΔN-Drosha does not grip the ssRNA-dsRNA junction as tightly as the full-length CED, which compromises the processing activity.

Pri-miRNAs contain several key structural features that facilitate the recognition by the Microprocessor and Drosha cleavage efficiency, including a 5’-UG-3’ motif at the basal junction (BJ) of the pri-miRNA hairpin, a 5’-UGU-3’/5’-GUG-3’ motif in the apical loop, and a 5’-CNNC-3’ motif downstream of the hairpin in the single-stranded flanking region positioned 16-18 nt from the Drosha cleavage site (Auyeung et al., 2013; Fang and Bartel, 2015). However, 5’-UGU-3’ or 5’-GUG-3’ motifs do not exist in miR-183/96/82 hairpins. The length of the stem of miR-182 and miR-183 hairpins is 48 bp and 52 bp, respectively, which are longer than 35±1 bp preferred by Drosha cleavage (Auyeung *et al*., 2013; Fang and Bartel, 2015). We speculate that the atypical sequence/structure of miR-183 and −182 hairpins is a critical determinant of Ago2-dependent cropping of pri-miR-183 but no other pri-miRNAs in ΔN-Drosha cells.

The nuclear localization of Drosha is mediated by a predicted nuclear localization signal (NLS) in the R/S-rich region(Lee *et al*., 2006). An alternatively spliced form of Drosha missing exon 6, which encodes NLS, localizes both in the nucleus and the cytoplasm (Dai et al., 2016; Link et al., 2016). Furthermore, the phosphorylation of serine (Ser)-300 or Ser-302 residue by glycogen synthase kinase 3β (GSK3β) is required for the nuclear retention of Drosha (Tang et al., 2011; Tang et al., 2010). We previously showed that p38 MAPK-dependent phosphorylation of Drosha at Ser-355 contributes to the nuclear-to-cytoplasmic shuttling of Drosha upon nutrients starvation (Jiang et al., 2021a). Because ΔN-Drosha is missing the NLS, Ser-300, Ser-302 and Ser-355, we predicted that ΔN-Drosha localizes exclusively in the cytoplasm; however, our results indicate that ∼70% of ΔN-Drosha is localized in the nucleus, indicating the presence of additional NLS or nuclear retention signals in aa 396-1374 of Drosha. Despite being expressed from the same loci and the mRNA amount being similar between ΔN-Drosha and FL, we noted the higher protein amount of ΔN-Drosha than FL. We previously reported that Drosha is degraded upon ubiquitination by Nedd4 (Jiang *et al*., 2021a). Ubiquitination of Drosha by Nedd4 requires the PPGY motif at aa 169-172 located in the NTR of Drosha, which binds the WW domains of Nedd4(Jiang *et al*., 2021a). Additionally, it has been reported that the stability of Drosha protein can be modulated by the acetylation of lysine (Lys)-48 in the NTR of Drosha by p300, CBP, and GCN5, which competes with ubiquitylation of the same Lys residues(Tang et al., 2013). Because ΔN-Drosha lacks both the PPGY motif and Lys-48, it would be expected to resist ubiquitin-proteasome-dependent degradation, leading to a higher accumulation than wild-type Drosha.

HHT is an autosomal dominant vascular disorder caused by the loss-of-expression or loss-of-function mutations in several mediators of the BMP signaling pathway, such as *Acvrl1, Endoglin*, and *Smad4 (Abdalla and Letarte, 2006)*. Missense mutations in NTR of Drosha, such as P32L, P100L, K226E, and R279L, were identified in individuals with HHT (Jiang *et al*., 2018). HHT patients with *Drosha* mutations exhibit a range of vascular defects stemming from abnormal vascular endothelial cell functions, such as epistaxis, telangiectasias, and arteriovenous malformations (AVMs)(Jiang *et al*., 2018). Like the ΔN protein, Drosha with P100L or R279L mutations fail to associate with Smad proteins and is unable to mediate BMP4-Smad1/5/8 dependent induction of miR-21 and miR-199a (Jiang *et al*., 2018). Based on our finding that ΔN-Drosha is unable to control RP biogenesis, protein synthesis, and cell proliferation upon nutrients deprivation, we speculate that the vascular phenotypes in HHT patients with mutant Drosha may be mediated by a combination of the loss of induction of BMP-dependent miRNAs and the loss of the vascular endothelial cell adaptation to changes in the extracellular environment.

We found that Ago2 processes the hairpins in pri-miR-183 in the absence of Drosha, or when Drosha is unable to process pri-miRNAs. Dicer-independent, Ago2-dependent cleavage of pre-miR-451 has been reported previously (Kretov *et al*., 2020; Yang and Lai, 2010; Yang *et al*., 2010). Ago2 is less efficient than Dicer in cleaving pre-miR-451 (Kretov *et al*., 2020). We found that Ago2 is more efficient than Drosha in processing pri-miR-183, because (i) miR-183/96/182 levels are higher in ΔN-Drosha than FL cells, and (ii) when Drosha is depleted in FL cells, miR-183/96/182 levels increase. An increased expression of miR-183 is reported in Wilms tumors, in which the catalytic activity of Drosha is frequently inactivated due mutations in the RNase III domains (Kruber *et al*., 2018; Ludwig et al., 2016; Rakheja *et al*., 2014; Spreafico *et al*., 2016; Torrezan *et al*., 2014; Walz *et al*., 2015; Wegert *et al*., 2015). Because Ago2 mutations have not been identified in Wilms tumors, we speculate that Ago2-dependent processing of pri-miR-183 might be facilitating the elevated miR-183 synthesis in Wilms tumors.

Although the majority of the miR-183 cluster miRNAs was identical to the reference sequence, ΔN-Drosha cells also contained miR-183 and miR-96 isomiRs that were 1- or 2-nt longer at the 5’-end. This suggests that the Ago2 cleavage site is less specific than the Drosha cleavage site. RNA-seq data showed that the levels of validated miR-183 target mRNAs, such as *ITGB1* (Li et al., 2010), *EZR* (Wang et al., 2008), *FOXO1* (McLoughlin et al., 2014), *RNF217* (Zhang et al., 2020), *PER2* (Zhou et al., 2021), *AXIN2* (Chen et al., 2018), and *MAL2* (Dambal et al., 2015), were higher in ΔN-Drosha cells than in FL cells, despite ΔN-Drosha cells expressing a higher amount of miR-183 than FL cells do (data not shown). We speculate that the addition of extra nucleotides at the 5’-end of isomiR-183 alters the seed sequence, resulting in the reduced recognition and/or silencing of miR-183 targets in ΔN-Drosha cells. The miR-183 cluster is abundantly expressed in the retina and plays an essential role in its development and homeostasis(Lumayag et al., 2013). Inactivation of individual or multiple miRNAs in the miR-183 cluster in mice leads to retinal degeneration (Lumayag *et al*., 2013; Wu et al., 2019; Xiang et al., 2017; Zhang *et al*., 2020). It has been reported previously that Ago2 is present in both retinal neurons and glia(Chu-Tan *et al*., 2021) and the depletion of Ago2 results in the depletion of miR-182 and miR-183 (Chen et al., 2021). Furthermore, a HITS-CLIP assay using whole retina lysates find that the miR-183 cluster is the most abundant group of miRNAs bound to Ago2 (Chu-Tan *et al*., 2021), which is consistent with our finding of the Ago2 role in pri-miR-183 processing. In the retina, depletion of Ago2 results in the reduction of miR-183 and −182 and retina degeneration(Chen *et al*., 2021) despite the presence of Drosha, suggesting that Ago2-mediated pri-miR-183 processing might be a predominant mechanism in the retina, which implicates a tissue-specific role of noncanonical processing like Ago2-dependent pre-miR-451 processing in erythrocytes(Kretov *et al*., 2020). Increased expression of the miR-183 cluster is associated with various human disorders, including cancer, autoimmune diseases, and neuronal diseases (Dambal *et al*., 2015). Furthermore, an increase in Ago2 protein amount along with post-translational modifications (PTMs) of Ago2, such as phosphorylation of tyrosine and serine and acetylation of Lys residues, are associated with poor prognosis and survival of cancer patients(Ye et al., 2015; Zhang et al., 2019). Our work suggests the induction of the miR-183 cluster is caused by the increased stability, cleavage activity, and/or cytoplasmic-to-nuclear translocation of Ago2, which then mediates pathogenesis (Dambal *et al*., 2015).

## Acknowledgement

We thank Drs. Brenton Graveley (Univ. of Connecticut) and Peng Du (Peking Univ.) for Lenti-Crispr-Drosha and -NS construct and IVP protocol, respectively. We also thank members of Hata lab for critical discussion.

## Author contributions

X.J. and A.H. conceived the project, designed experiments, analyzed data, and drafted the manuscript. A.P., S.H., J.T., P.G. and G.L. performed experiments, and edited the manuscript.

## Methods

### Cell culture

Human embryonic kidney (HEK) 293T cells (ATCC, CRL-3216) were cultured in DMEM (high glucose, Sigma-Aldrich, SH30022.01) with 10% FCS (Hyclone, SH3007103), 1% penicillin/streptomycin at 37 °C and 5% CO_2_. Drosha ΔN-Drosha clones were maintained in the same media except containing 5 nM puromycin. For serum starvation, cells were cultured in DMEM with 1% FCS. To downregulate the *Drosha, Ago2*, and *Dgcr8* expression in FL-Drosha and ΔN-Drosha clones, siRNAs were introduced into clones using LipofectamineTM RNAiMAX transfection Reagent (Invitrogen, 13778-150).

### Generation of Drosha ΔN-Drosha cell lines by CRISPR-Cas9 genome editing

Two gRNAs were designed for each targeted human Drosha gene to facilitate genomic deletion of exon 5 using several online tools. The sequence of two gRNAs is listed in Supplementary information. gRNAs were cloned in the lentiCRISPR v2 plasmid (Addgene plasmid #52961). HEK293T cells were transfected simultaneously with PMD2.G (Addgene plasmid #12259), and psPAX2 (Addgene plasmid #12260) by lipofectamine2000 (Invitrogen, 11668030) to generate the lentivirus. Six h after the transfection, the culture media were replaced with DMEM (high glucose, Sigma-Aldrich) with 10% FCS. After 48 hr, supernatant was collected, filtered with 0.45μm filter, and used to infect HEK293T cells with polybrene (8μg/ml, Sigma-Aldrich, TR-1003). The infected HEK293T cells were selected in the media containing puromycin (5ng/μl). After the puromycin selection, growing cells were trypsinized, and single cells were sorted into 96-well plates by FACS (FACS AriaIII, BD Biosciences). After 30 days, clonal cell lines were expanded and subjected to genotyping. The genotyping primer sequence are listed in Supplementary information.

### Identification of Drosha ΔN-Drosha clones by genomic PCR

To distinguish the wild type Drosha alleles from the mutant (ΔN-Drosha) alleles, trypsinized cells were pelleted, resuspended in 10 mM Tris (pH 8.7), heated at 95°C for 10 min, incubated with proteinase K (0.5 µg/µl) for 20 min at 37°C, inactivated at 95°C for 15 min, and used as DNA template for genomic PCR analysis. The condition of genomic PCR is as follows: initial denaturing reaction at 95°C for 2 min; 35 cycles of denaturation at 95°C for 30 sec, annealing at 56°C for 30sec, and elongation at 72°C for 30 sec, followed by the final extension reaction at 72°C for 5 min. Primers designed to amplify the genomic region surrounding the site of deletion (primer #1-3) are listed in Supplementary information. The wild type allele yields no PCR fragments by primer #1 and #2 but yield PCR fragments (621 bp) by primer #1 and #3. The exon 5-deleted (Δex5) allele yields PCR fragments (530 bp) by primer #1 and #2 but no fragments by primer #1 and #3.

### SDS-PAGE and Immunoblot analysis

Cell lysates were denatured in sample buffer (Invitrogen, NP0007) with reducing agent (Invitrogen, NP0009) for 5 min at 95°C, separated by SDS-gel electrophoresis, and transferred to nitrocellulose blotting membrane (Genesee Scientific). Membranes were blocked with TBST (20 mM Tris, 150 mM NaCl, pH 7.6, and 0.1% Tween20) with 3% nonfat dry milk for 1 h and incubated with primary antibody in TBST with 1% milk overnight at 4°C. Chemiluminescence signals were detected using SuperSignal™ West Dura extended duration substrate (ThermoFisher, 34076) and imaged using an Odyssey Dlx Imaging System (LI-COR). Following antibodies were used for immunoblot: Anti-Drosha antibody (1:500 dilution, Bethyl, A301-866A), anti-Dgcr8 (1:500 dilution, Proteintech,10996-1-AP), anti-Ddx5 (1:200 dilution, Abcam, ab21696), anti-Ago2 (1:500 dilution, Cell signaling Technology, 2897), anti-GAPDH (1:5000 dilution, Millipore, MAB374), anti-Lamin A/C (1:2500 dilution, Cell signaling Technology, 2032), anti-Gata1 (1:200 dilution, R&D, MAB17791-SP), anti-puromycin (1:2000 dilution, Kerafast, 3RH11), anti-Rpl11 (1:300 dilution, Proteintech,16277-1-AP), anti-Rpsa (1:300 dilution, Abcam, ab137388), anti-Rps24 (1:300 dilution, Abcam, ab102986), anti-Rps26 (1:300 dilution, Abcam, ab104050), anti-Rps19 (1:300 dilution, Santa Cruz Biotechnology, sc-100836), anti-Smad1 (1:500 dilution, Invitrogen, 38-5400), phospho-Smad1/5/8 (1:100 dilution, Cell signaling Technology, 9511), anti-β-actin (1:5000 dilution, Sigma-Aldrich, A5441), IRDye 680RD goat anti-rabbit IgG (H + L) (LI-COR, 926-68071), IRDye 800CW goat anti-rabbit IgG (H + L) (LI-COR, 926-32211), IRDye 680RD goat anti-mouse IgG (H + L) (LI-COR, 926-68070), and IRDye 800CW goat anti-mouse IgG (H + L) (LI-COR, 926-32210). anti-Rabbit-IgG-HRP-linked (1:3000 dilution, Cell signaling Technology, 7074), anti-Mouse-IgG-HRP-linked (1:3000 dilution, Cell signaling Technology, 7076),anti-Rabbit-IgG-HRP-linked (1:3000 dilution, Cell signaling Technology, 7077).

### Quantitative reverse transcriptase-polymerase chain reaction (qRT-PCR) analysis

Total RNAs were extracted by RNeasy Mini Kit (#74104, Qiagen) and subjected to generate cDNAs by RT reaction (#17088890, Bio-Rad). qPCR was performed using iQ SYBR Green Supermix (#1708882, Bio-Rad). All reactions were run in triplicates. The relative expression values were determined by normalization to *GAPDH* transcript levels and calculated using the ΔΔCT method. Primers used for qRT-PCR are listed in Supplementary information.

### Quantitative miRNA analysis

Mature miRNAs were determined using TaqMan microRNA Assays (Applied Biosystems Inc.). Normalization was performed with the small nuclear RNA U6 (RNU6B; Applied Biosystems Inc.). All real-time reactions, including no-template controls and real-time minus controls, were run using the CFX Connect Real-Time PCR System (Bio-Rad) and performed in triplicate. Relative expression was calculated using the ΔΔCT method.

### Immunoprecipitation assay

Cells were lysed in SBB buffer (1% Triton X-100, 150mM NaCl, 50mM Tris-Cl at pH 7.5, 1mM EDTA) supplemented with protease inhibitors (Sigma, P8340) and phosphatase inhibitor (Sigma, P5726). Cell lysates were incubated at 4°C for 30 min and centrifuged at 12,000 g for 10 min at 4°C. Lysates were incubated with anti-Drosha, anti-Ago2, anti-Dgcr8, and anti-IgG (negative control) nutating overnight at 4°C followed by the addition of Dynabeads™ Protein A/G (Invitrogen, 10002D/10004D) and rocking for 4 h at 4°C. The magnetic beads were precipitated and rinsed with SBB buffer for 5 min at 4°C for three times, followed by adding sample buffer (Invitrogen, NP0007) with reducing agent (Invitrogen, NP0009) and heated at 95°C for 3 min.

### Nuclear and cytoplasmic fractionation

Cells were washed with PBS twice, scrape off and pelleted by centrifuging at 4,500 g for 5 min. Cells were then swelled by adding 5 volume of lysis buffer (10 mM HEPES, pH 7.9, with 1.5 mM MgCl2, 10 mM KCl, 1mM DTT and protease inhibitor, sigma, P8340) and homogenized. After centrifugation at 10,000 g for 15 min, the supernatant was collected as a cytoplasmic fraction. The crude nuclei pellet was resuspended in 2/3 volume extraction buffer (20 mM HEPES, pH 7.9, with 1.5 mM MgCl2, 0.42 M NaCl, 0.2 mM EDTA, 25% (v/v) Glycerol, 1mM DTT and protease inhibitor sigma P8340) and homogenized with a tissue homogenizer. After centrifuging at 20,000 g for 5 min, the supernatant was collected as a nuclear fraction.

### Next generation RNA-seq and small RNA-seq and analysis

Total RNAs were extracted from cells using TRIzol (Invitrogen). The quality of RNAs was evaluated by 2100 Bioanalyzer Instrument (Agilent Technologies) and the samples with RIN>8.0 were sent to Beijing Genome Institute (BGI) for RNA sequencing and small RNA sequencing. The sequencing was performed with DNBSEQ™ technology platforms. Adapter removed clean data were generated by BGI. Quality control, index generation and mapping for RNA sequencing were done with Salmon software tool. Differential gene expression was analyzed with R package DESeq2_1.4.5. Quality control for small RNA sequencing data were done by fast quality filter and fastx trimmer. Index file was built by bowtie 2 with GRCH38. The subsequent mapping was done with miRDeep2.

### Chromatin Immunoprecipitation (ChIP) assay

Cells were crosslinked treated with 1% Formaldehyde for 15 min at room temperature Followed by quenching with 1M Glycine, cells were washed with PBS and lysed with lysis buffer (50 mM Tris-Cl pH 8.1, 10 mM EDTA, 1% SDS and protease inhibitor). Genomic DNAs were sheared to average length of 200-500bp by sonication, followed by clearing lysates by centrifugation at 12,000g for 10 min at 4°C. Incubate the supernatant with protein A/G dynabeads (invitrogen 10002D/10004D) for 1 h at 4°C, dilute the pre-cleared sample to 1:10 ration with dilution buffer (20 mM Tris-Cl pH 8.1, 150 mM NaCl, 2 mM EDTA, 1% Triton X-100 and protease inhibitor) and 1/10 volume was kept as input before incubation with anti-Drosha overnight at 4°C followed by 1 h incubation with protein A/G dynabeads at 4°C. After the dynabeads were washed with a buffer I (20 mM Tris-Cl pH 8.1, 150 mM NaCl, 2 mM EDTA, 1% Triton X-100, 0.1% SDS), buffer II (20 mM Tris-Cl pH 8.1, 500 mM NaCl, 2 mM EDTA, 1% Triton X-100, 0.1% SDS), and buffer III (10mM Tris-Cl pH8.1, 250mM LiCl, 1mM EDTA, 1%NP-40, 1% Deoxycholate) at 4°C, the dynabeads were further washed twice with cold TE (10 mM Tris-Cl pH 8.1, 1 mM EDTA). The dynabeads were incubated in 250 μl elution buffer (200 mM NaHCO_3_, 1% SDS) at room temperature for 15 min twice. The eluates were mixed with 1/25 volume 5M NaCl and incubated at 65°C for 4 h. 1/50 volume of 0.5 M EDTA, 1/25 volume of Tris-Cl pH 6.5, 1/100 volume of Proteinase K (10 mg/ml) were added and incubated at 45°C for 1 h. Precipitated genome fragments were purified with QIAquick PCR Purification Kit, followed by PCR analysis. Primers used for ChIP assay are listed in Supplementary information.

### Proliferation assay

Cell growth was monitored by cell counting and 3-(4,5-dimethylthiazol-2-yl)-2,5-diphenyltetrazolium bromide (MTT) assay using MTT cell growth assay kit (#CT02, Millipore). FL or ΔN-Drosha cells (1×10^5^) were seeded in 12-well plates and cultured in DMEM containing 10% or 1% FCS. 8, 16, 24, 32, or 40 h after the media change, cells were harvested and counted by a hemocytometer. For MTT assay, MTT dye was added to each well, incubated at 37°C for 1 h, followed by the addition of 0.1 mL isopropanol with 0.04 N HCl. The absorbance was measured at a wavelength of 570 nm.

### Puromycin incorporation assay

Cells were cultured in the growth media (DMEM with 10% FCS) or starvation media (DMEM with 1% FCS) for 16 h, followed by the treatment with 1μM puromycin at 37°C for 10 min. Total cell lysates were generated and subjected to SDS-PAGE and immunoblot with an anti-puromycin antibody (Kerafast, EQ0001).

### In vitro pri-miRNA processing (IVP) assay

A partial pri-let-7b (432-nt) or Pri-miR-183 (327-nt) sequence was amplified from human genomic DNAs and used as *in vitro* transcription template in the reaction using Riboprobe System-T7 Kit (P1440, Promega) in the presence of 0.4μL 5-(3-Aminoallyl)-uridine-5’-triphosphate labeled with ATTO 680(Aminoallyl-UTP-ATTO-680, NU-821-680, Jena bioscience) in each reaction. Cells from 10-cm dish were harvested in 300μl sonication buffer [20 mM Tris-HCL pH=8.0, 100 mM KCL, 0.2 mM EDTA RNase-free] and sonicated with 20% intensity for 5 sec for 3 times. After the sonication, cell lysates were subjected to the centrifugation (12,000 rpm at 4 °C for 15 mins). IR-680 labelled pri-let-7b or pri-miR-183 (0.1 μg) was mixed with the supernatant (total protein amount of 20-30 μg) supplemented with 6.4 mM MgCl_2_ and 0.5 U/μl Recombinant RNase Inhibitor (Promega) in total volume of 30 μl. After the incubation at 37°C for 30 min, the reaction mixtures were separated on a 15% Urea-PAGE gel at 90 V for 150 min to separate the processing product [pre-let-7b (82-nt) or pre-miR-183 (110-nt)] from the substrate (pri-let-7b or pri-miR-183). The gel image was captured by Odyssey Dlx Imaging System (LI-COR). The processing activity was quantitated by the amount of pre-miRNA divided by the total amount of pri-miRNA and pre-miRNA.

### RNA Immunoprecipitation (RIP) assay

HEK293T cells with FL and ΔN-Drosha Drosha were subjected to crosslinking with 1% formaldehyde for 15 min at room temperature. Nuclei were isolated and disrupted by sonication using Bioruptor (Diagenode). The sonicated lysates were cleared and subjected to immunoprecipitation with anti-Ago2, anti-Drosha, and anti-Dgcr8. After immunoprecipitation, washing and elution, the cell pellets were subjected to 10 U DNase I treatment for 30 min at 37 °C to remove any remaining DNA. Next, RNA was extracted using Trizol/phenol:chloroform (5:1), precipitated with ethanol, and dissolved in 20ul of nuclease free water. 5ul of RNA was used for 20ul cDNA synthesis reaction. qRT-PCR reactions were performed using pre-miR-primers by real-time PCR machine (CFX96, BioRad). 100 U/ml RNase inhibitor (SUPERase•in™) was used throughout the experiment.

### Statistical analysis

Graphs were generated with GraphPad PRISM software. Statistical significance was calculated in R version 3.2.3 by Student’s t test. The null hypothesis of the medians/means being equal was rejected at α = 0.05 and p values were generated by unpaired Student’s t test and presented in figures. The sample size was estimated by power analysis and is presented in the figure legend. All experiments were performed at least three times with biological triplicates each time.

### Data availability

RNA sequencing data are under submission and will be available at the NCBI Sequence Read Archive (SRA) shortly [GEO Accession Number (TBA)].

### Lead Contact

Further information and requests for resources and reagents should be directed to and will be fulfilled by the Lead Contact, Akiko Hata.

