## Supplementary Figs and information for "Essential role of the amino-terminal region of Drosha for the Microprocessor function"

Supplementary Fig. S1 Prabhakar et al.

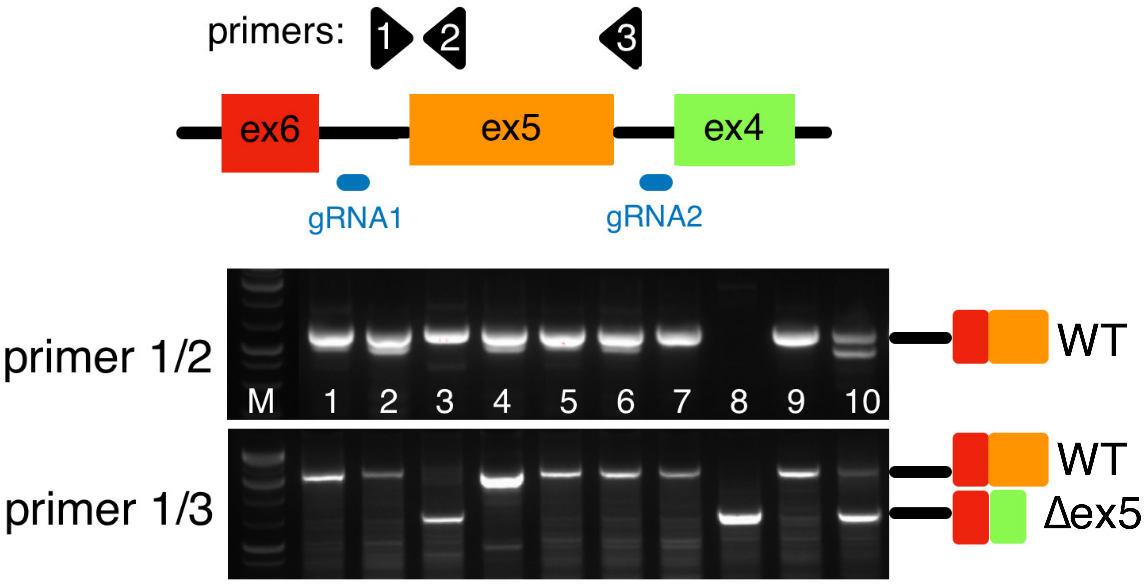


**Supple. Fig. S1 Identification of *Drosha* mutant clones in HEK293T cells.** Two guideRNAs (gRNA1 and gRNA2, blue) were used to delete exon5 (Δex5) of *Drosha* by CRISPR/Cas9-based genome editing are indicated. Lanes 1-10 correspond to the genomic DNA from clones 1-10. Primers 1-3 (top) were used for genomic DNA analysis to distinguish wild type (WT) vs Δex5 allele of the Drosha gene. Results of PCR analyses of genomic DNA by 2 set of primers are shown (top: primers #1 and #2 and bottom: primers #1 and #3). M: molecular marker.

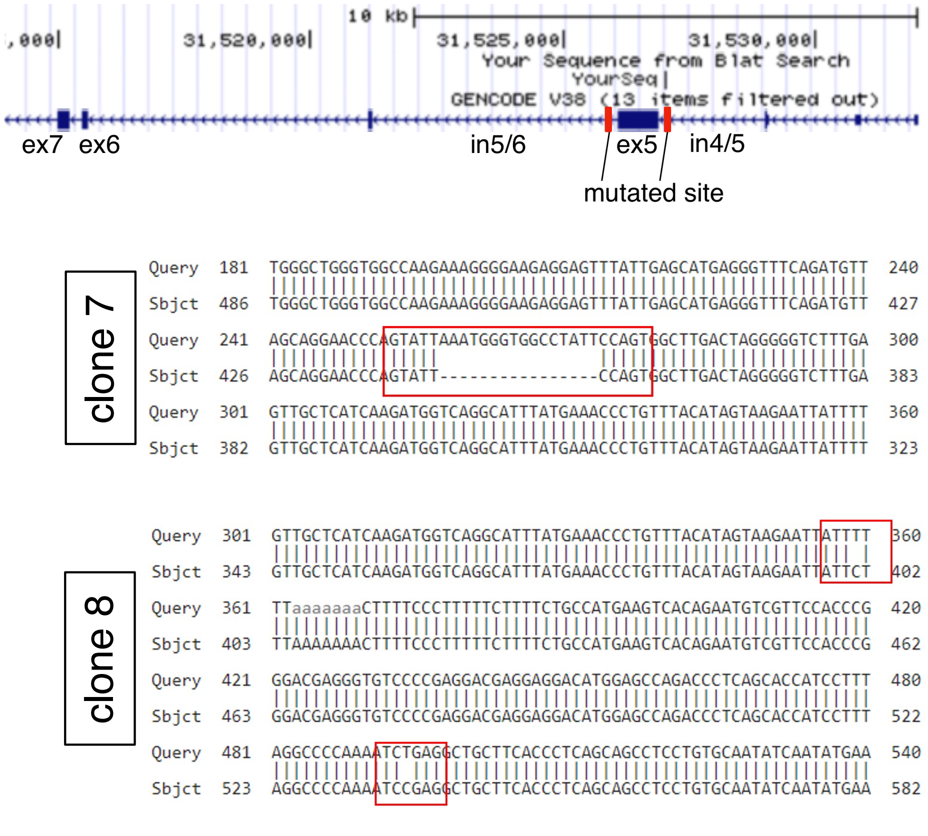


**Supple. Fig. S2 Mutations upstream of exon5 of the Drosha gene were identified in clone 7 and 8.** Two gRNAs were used to delete a whole exon 5 (ex5) of human *Drosha* by CRISPR/Cas9-based genome editing (top). Mutated sequence in intron 4/5 in clone 7 (Δex5/+) and clone 8 (Δex5/Δex5) are shown (bottom).

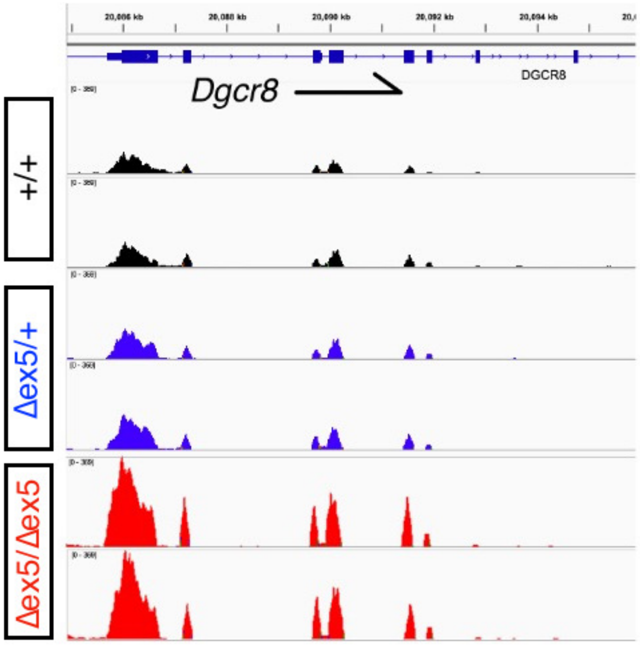


**Supple. Fig. S3 RNAseq analysis of the Dgcr8 transcripts in clone 4 (+/+), 7 (Δex5/+), and 8 (Δex5/Δex5)** RNA-seq analysis confirms an increased level of reads corresponding to the Dgcr8 mRNA in Δex5/Δex5 cells (clone 8, red) in comparison with +/+ cells (clone 4, black) and Δex5/+ cells (clone 7, blue). Two libraries were generated from each clone and both RNA-seq data are shown.

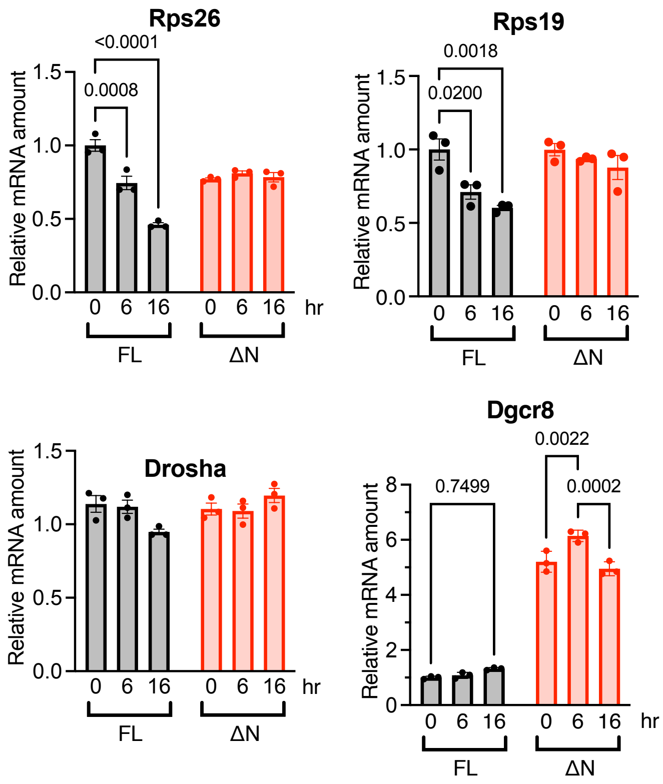


**Supple Fig. S4 No reduction of the RP mRNAs in ΔN-Drosha cells under serum starvation** qRT-PCR analysis of the Rps19, Rps24, Drosha and Dgcr8 mRNAs relative to GAPDH mRNAs in FL (+/+) cells (black) and ΔN-Drosha (Δex5/Δex5) cells (red). Results are plotted as mean± SEM. n=3 independent experiments.

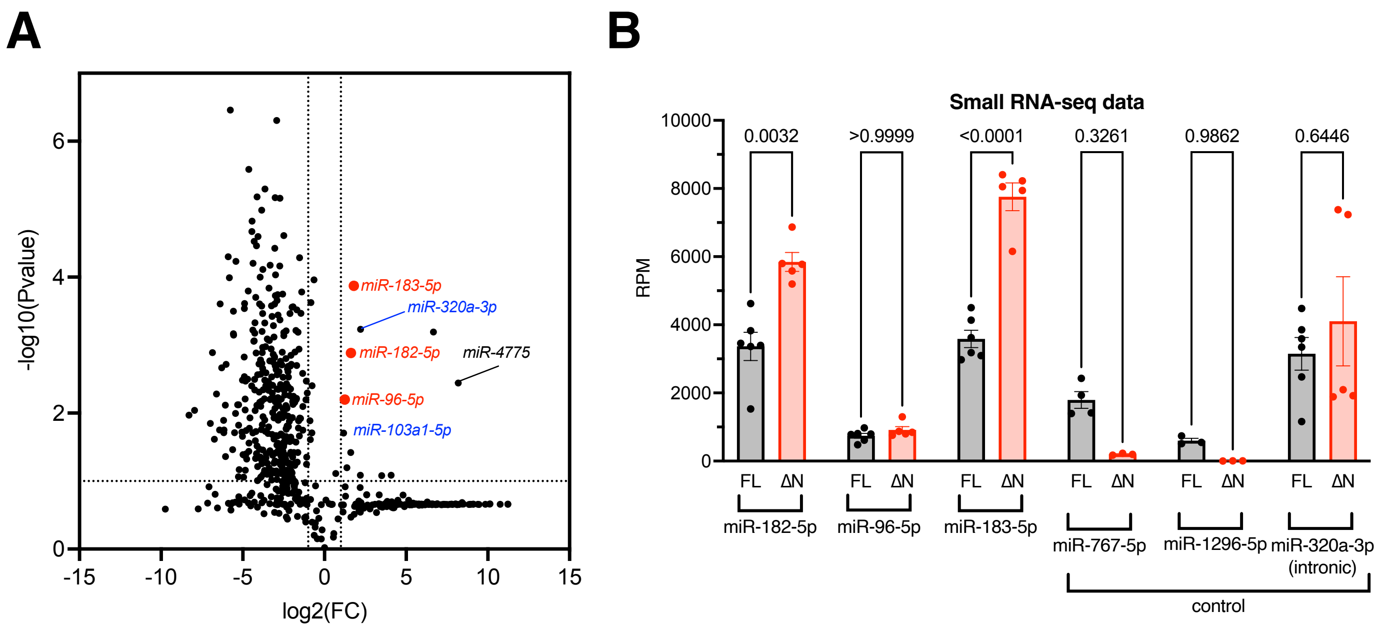


**Supple Fig. S5 Higher amount of miR-183 cluster in ΔN-Drosha cells compared to FL-Drosha cells A.** Global analysis of miRNA expression in FL (+/+) and ΔN-Drosha (Δex5/Δex5) cells is visualized in volcano plot. The fold change (FC) of miRNAs ($log_2$FC) are shown in X axis and p-values [-log10(P-value)]in Y-axis. miR-183 cluster of miRNAs: miR-182, miR-96, and miR-183 are shown in red. Intronic miRNAs whose processing is Drosha-independent are shown in blue. **B.** Small RNA-seq data of FL cells and ΔN-Drosha cells indicates similar level of expression of miR-182-5p, miR-96-5p and miR-183-5p in clones 4 and 8 despite miR-767-5p and miR-1296-5p (control) were significantly less in clone 8. A similar level of intronic miR-320a-3p was detected in FL and ΔN cells. Results are plotted as mean RPM (read per million reads)± SEM. n=6 for FL cells and n=5 for ΔN-Drosha cells.

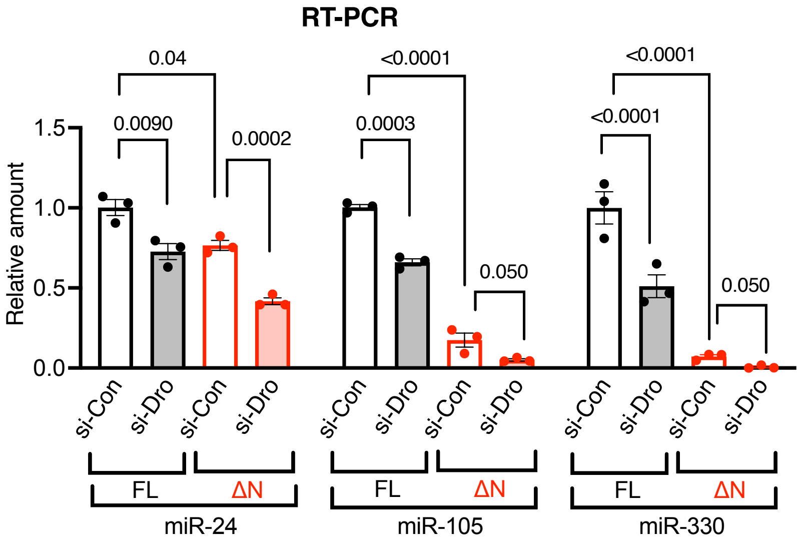


**Supple. Fig. S6 Drosha-dependent processing of miR-24, -105 and -330** FL or ΔN-Drosha cells transfected with siRNA against Drosha (si-Dro) or non-specific control siRNA (si-Con) were subjected to qRT-PCR analysis of miR-24, -105, and -330. Relative amount of miRNAs normalized to U6 snRNA is plotted as mean± SEM. n=3.

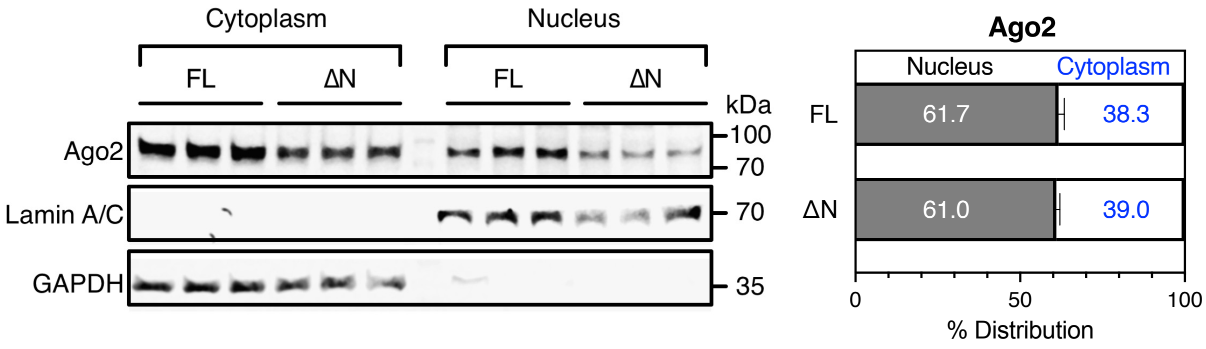


**Supple. Fig. S7 Ago2 localizes not only in the cytoplasm but in the nucleus** Nuclear and cytoplasmic fraction of FL and ΔN-Drosha cells were subjected to immunoblot analysis of Ago2, Lamin A/C (control for the nucleus), and GAPDH (control for the cytoplasm) (left). Relative distribution (%) of FL and ΔN-Drosha in the nucleus vs cytoplasm is shown (right).

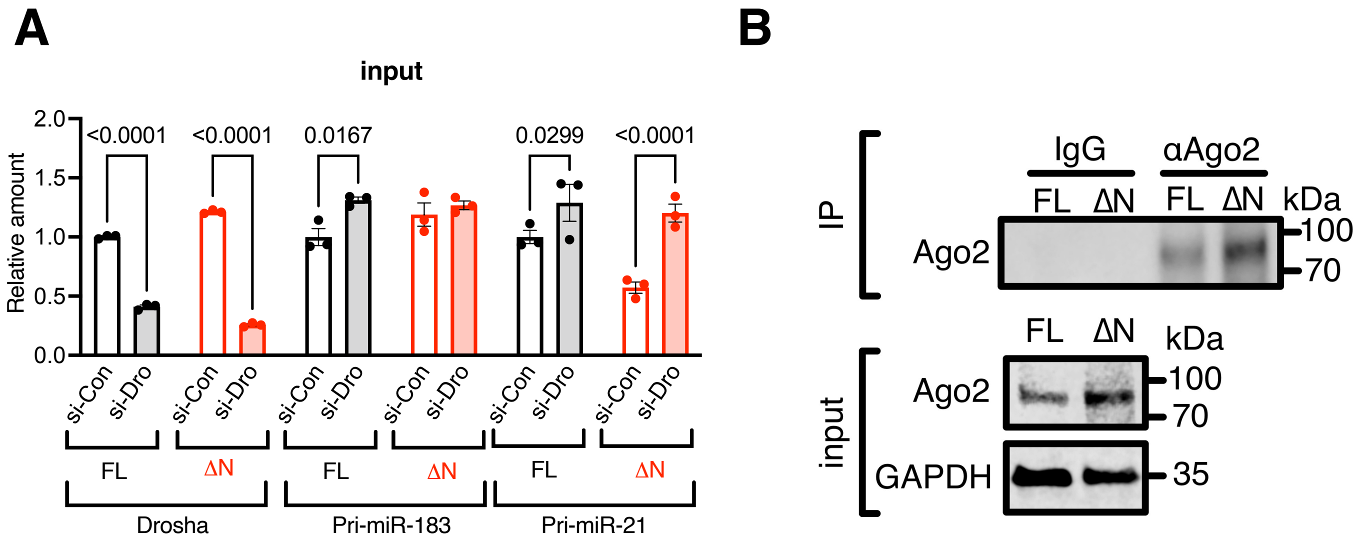


**Supple. Fig. S8 Similar amount of Ago2 protein was precipitated by anti-Ago2 antibody. A.** The relative amount of Drosha, pri-miR-183, and pri-miR-21 mRNAs in FL and ΔN-Drosha cells normalized to GAPDH mRNA in the input samples of RIP assay (anti-Drosha IP) is plotted as mean± SEM. n=3 **B.** IP with anti-Ago2 antibody (αAgo2) but not nonspecific IgG (IgG) pulls down Ago2 protein together with associating RNAs in FL and ΔN-Drosha cells (top, IP).

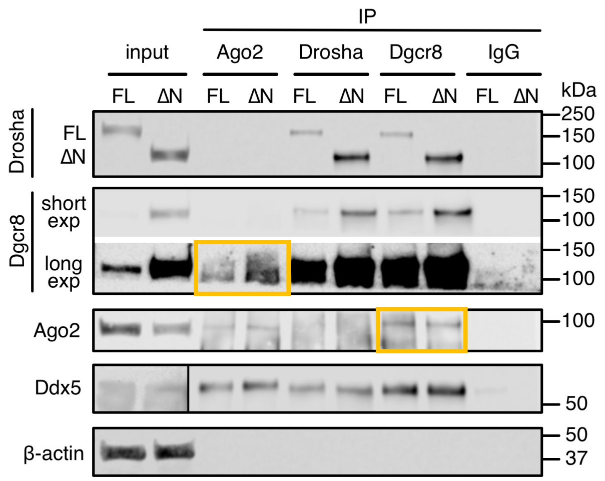


**Supple. Fig. S9 Association of Dgcr8 with Ago2.** Ago2, Drosha, Dgcr8 were immunoprecipitated in FL and ΔN-Drosha cells, followed by immunoblot with anti-Drosha, anti-Dgcr8, anti-Ago2, or anti-Ddx5 antibody. Interaction between Ago2 and Dgcr8 were detected in both Ago2 IP and Dgcr8 IP samples (yellow boxes). Non-specific IgG was used as control IP. Anti-β-actin antibody was used as loading control for the immunoblot.

### Supplementary information

**1. A list of PCR primers:** Sequences of the PCR primers for genomic PCR amplification, RT-PCR, miRNA, RIP, IVP, and ChIP are listed below. All primers are for human.

| **Primer Name** | **Primer Sequence** | **Annotation** |
|---|---|---|
| *DROSHA*-in3-F (primer #1) | 5'-GCATTTGGAGATGGGAGTG-3' | Genotyping of Drosha locus |
| *DROSHA*-in4-R (primer #2) | 5'-GGGCAACATAGCGAGATTC-3' | |
| *DROSHA*-ex5-R (primer #3) | 5'-GGAAGGGTACAAAGTCTGGTCG-3' | |
| gRNA1 | 5'-GGAACCCAGTATTAAATGGGTGG-3' | guideRNAs for Crispr-Cas9 Drosha gene editing |
| gRNA2 | 5'-CCAAACCAAAAGGATCCAGTAGG-3' | |
| *DROSHA*-ex3-qPCR-F | 5'-ACATATCCAGGCGGAACATC-3' | qRT-PCR for Drosha ΔN-Drosha |
| *DROSHA*-ex5-qPCR-R | 5'-ATGGTGATCTTCGGTTGTCTC-3' | |
| *DROSHA*-qPCR-F | 5'- GAAACTTCGCCACCTCCTAGCA-3' | qRT-PCR for Drosha WT |
| *DROSHA*-qPCR-R | 5'- CTCCACCGTTACTTCTCGTCTC-3' | |
| *DGCR8*-qPCR-F | 5'- GGTCCGCCCTGTCTATAATTTC-3' | qRT-PCR for Dgcr8 |
| *DGCR8*-qPCR-R | 5'- GAGTCCTCGATGCTGATGTG-3' | |
| *Rps19-qPCR-F* | 5'- ACTTCAGCCGAGGCTCCAAGAG-3' | qRT-PCR for Rps19 |
| *Rps19-qPCR-R* | 5'- CTCTTTGTCCCTGAGGTGTCAG-3' | |
| *Rps24-qPCR-F* | 5'- CCAATGTTGGTGCTGGCAAAAAG-3' | qRT-PCR for Rps24 |
| *Rps24-qPCR-R* | 5'- GCACTTCTACCTGCCACACAAC-3' | |
| *Rps26-qPCR-F* | 5'- GGACATTTCTGAAGCGAGCGTC-3' | qRT-PCR for Rps26 |
| *Rps26-qPCR-R* | 5'- CGATTCCTGACTACTTTGCTGTG-3' | |
| *Rps2-chip-F* | 5'- GCCACACTGACTAGTTCCTTC-3' | ChIP for Rps2 locus |
| *Rps2-chip-R* | 5'- CCACTGCCGAAACCTCC -3' | |
| *Rps10-chip-F* | 5'- GTTCCATCGGCTCCCATC-3' | ChIP for Rps10 locus |
| *Rps10-chip-R* | 5'- CCCCTACCCCATAAAATAAGCC-3' | |
| *Rpl28-chip-F* | 5'- TTTTCCCCTCACTCTCATTCG-3' | ChIP for Rpl28 locus |

| | | |
|---|---|---|
| *Rpl28-chip-R* | 5’- ACTGGGAACTTGGGTGAATG-3’ | |
| *pre-miR-21-F* | 5’- TGTCTGCTTGTTTTGCCT-3’ | qRT-PCR and RIP assay for miR-21 locus |
| *pre-miR-21-R* | 5’- GGATATGGATGGTCAGATGAA-3’ | |
| *pre-miR-199a-F* | 5’- GCCAACCCAGTGTTCAGACTA-3’ | qRT-PCR for miR-199a |
| *pre-miR-199a-R* | 5’- GCCTAACCAATGTGCAGACTA-3’ | |
| *pri-miR-183/96/182-F* | 5’- TGAAGGGGAACATTGGCCTC-3’ | qRT-PCR and RIP assay for miR-183 cluster locus |
| *pri-miR-183/96/182-R* | 5’- GGTCATCTCCGAACAGCTCC-3’ | |
| *pri-let-7b#1-F* | 5’-TAATACGACTCACTATAGGGACTTCCCAAGACCAGCC-3’ | IVP assay for let-7b |
| *pri-let-7b#1-R* | 5’-GGGGCCAGTTCCAAGTTCATGG-3’ | |
| *pri-miR-183-F* | 5’-TAATACGACTCACTATAGCCGCAGAGTGTGACTCCTG-3’ | IVP assay for miR-183 |
| *pri-miR-183-R* | 5’-CACCCTTGGAAGCAGCCT-3’ | |
| *pri-let-7b#2-F* | 5’- CCCTACCTCAGTGACACGAC-3’ | RIP assay for let-7b locus |
| *pri-let7-b#2-R* | 5’- ATCTAGCTCCCAGATGCCCA-3’ | |

### 2. A list of reagents

| # | Reagent | Company | Catalog no. | |
|---|---|---|---|---|
| 01. | Puromycin | InvivoGen | ant-pr-1 | |
| 02. | DMEM-high glucose | HyClone lab | SH30022.01 | |
| 03. | Lipofectamine2000 | Invitrogen | 11668-030 | |
| 04. | Lipofectamine RNAiMax | Invitrogen | 13778-150 | |
| 05. | SuperSignal™ West Dura extended duration substrate | ThermoFisher | 34076 | |
| 06. | Nitrocellulose blotting membrane | Genesee Scientific | 84-875 | |
| 07. | Polybrene | Sigma-Aldrich | TR-1003 | |
| 08. | Trypsin | Life technologies | 25200-072 | |
| 09. | Proteinase K | Invitrogen | P/N100005393 | |
| 10. | Protease Inhibitor | Sigma | P8340 | |
| 11. | Phosphatase Inhibitor | Sigma | P5726 | |
| 12. | Riboprobe System-T7 Kit | Promega | P1440 | |
| 13. | ATTO 680 | Jena bioscience | NU-821-680 | |
| 14. | RNase Inhibitor | Promega | | IVP |
| 15. | RNase inhibitor | Invitrogen | AM2696 | RIP |
| 16. | DNase I | Ambion | AM2238 | |
| 17. | SDS-PAGE sample buffer | Invitrogen | NP0007 | |
| 18. | SDS-PAGE reducing agent | Invitrogen | NP0009 | |

### 3. A list of antibodies: Following antibodies were used for immunoprecipitation, immunoblot, ChIP, or RIP assay.

| # | Antibody | Company | Catalog no. |
|---|---|---|---|
| 01. | Ago2 | Cell signaling Technology | 2897 |

| | | | |
|---|---|---|---|
| 02. | Beta-actin | InvivoGen | ant-pr-1 |
| 03. | Ddx5 | Abcam | ab21696 |
| 04. | Dgcr8 | Proteintech | 10996-1-AP |
| 05. | Drosha | Bethyl | A301-866A |
| 06. | GAPDH | Millipore | MAB374 |
| 07. | Gata1 | R&D systems | MAB17791-SP |
| 08. | Lamin A/C | Cell signaling Technology | 2032 |
| 09. | Puromycin | Kerafast | 3RH11 |
| 10. | Rpl11 | Proteintech | 16277-1-AP |
| 11. | Rps19 | Santa Cruz Biotechnology | sc-100836 |
| 12. | Rps24 | Abcam | ab102986 |
| 13. | Rps26 | Abcam | ab104050 |
| 14. | Rpsa | Abcam | ab137388 |
| 15. | Smad1 | Invitrogen | 38-5400 |
| 16. | p-Smad1/5/8 | Cell signaling Technology | 9511 |
| 17. | IRDye-680RD goat anti-rabbit IgG (H+L) | Li-Cor | 926-68071 |
| 18. | IRDye-800CW goat anti-rabbit IgG (H+L) | Li-Cor | 926-32211 |
| 19. | IRDye-680RD goat anti-mouse IgG (H+L) | Li-Cor | 926-68070 |
| 20. | IRDye-800CW goat anti-mouse IgG (H+L) | Li-Cor | 926-32210 |
| 21. | anti-Rabbit-IgG-HRP-linked | Cell signaling Technology | 7074 |
| 22. | anti-Mouse-IgG-HRP-linked | Cell signaling Technology | 7076 |
| 23. | anti-Rabbit-IgG-HRP-linked | Cell signaling Technology | 7077 |

### 4. A list of miRNA quantitation reagents and siRNAs

| **#** | **miRNA** | **Company** | **Catalog no.** |
|---|---|---|---|
| 01. | hsa-miR-183 Taqman assay | Applied Biosystems | 4427975-002269 |
| 02. | has-miR-182 Taqman assay | Applied Biosystems | 4427975-002334 |
| 03. | hsa-miR-96 Taqman assay | Applied Biosystems | 4427975-000186 |
| 04. | hsa-miR-21 Taqman assay | Applied Biosystems | 4427975-000397 |
| 05. | hsa-miR-103 Taqman assay | Applied Biosystems | 4427975-000439 |
| 06. | hsa-miR-105 Taqman assay | Applied Biosystems | 4427975-002167 |
| 07. | hsa-miR-199a Taqman assay | Applied Biosystems | 4427975-000498 |
| 08. | hsa-miR-24 Taqman assay | Applied Biosystems | 4427975-000402 |
| 09. | hsa-miR-34a Taqman assay | Applied Biosystems | 4427975-000426 |
| 10. | hsa-miR-330 Taqman assay | Applied Biosystems | 4427975-002230 |
| 11. | siAgo2 | Sigma-Aldrich | SASI-Hs02_00343736 |
| 12. | siDgcr8 | Sigma-Aldrich | SASI-Hs02_00355944 |
| 13. | siDrosha | Dharmacon™ | L-016996-00-0005 |
| 14. | siControl | Dharmacon™ | D-001206-13-05 |
